# Direct serotonin release in humans shapes decision computations within aversive environments

**DOI:** 10.1101/2023.09.30.560111

**Authors:** Michael J Colwell, Hosana Tagomori, Fei Shang, Hoi Cheng, Chloe Wigg, Michael Browning, Phil J Cowen, Susannah E Murphy, Catherine J Harmer

**Author notes:** Joint Senior Authorship. Authors for Correspondence:* Catherine J Harmer, Michael J Colwell.

## Abstract

The role of serotonin in human behaviour is critically informed by approaches which allow *in vivo* modification of synaptic serotonin. However, characterising the effects of increased serotonin signalling in human models of behaviour is challenging given the limitations of available experimental probes (*e.g.*, SSRIs). Here we use a now accessible approach to directly increase synaptic serotonin in humans – a selective serotonin releasing agent – and examine its influence on domains of behaviour historically considered core functions of serotonin. Computational techniques including reinforcement learning and drift diffusion modelling were fit to observed behaviour. Reinforcement learning models revealed that increased synaptic serotonin reduced sensitivity specifically for outcomes in aversive but not appetitive contexts. Furthermore, increasing synaptic serotonin enhanced behavioural inhibition, and shifted bias towards impulse control during exposure to aversive emotional probes. These effects were seen in the context of overall improvements in memory for neutral verbal information. Our findings highlight the direct effects of increased synaptic serotonin on human behaviour, underlining its critical role in guiding decision-making within aversive and neutral contexts, and offering broad implications for longstanding theories of central serotonin function.

## Introduction

Understanding the function of central serotonin (or 5-hydroxytryptamine, 5-HT) has been a focal goal of neuroscience research for nearly a century ^1^, not least because of its central role in the effects of many psychiatric drugs, predominantly selective serotonin reuptake inhibitors [SSRIs], and street drugs (*e.g.,* ±3,4-methylenedioxymethamphetamine [MDMA] and lysergic acid diethylamide) ^2, 3^. Serotonin is phylogenetically ancient, and its function translates across species to many lower- and higher-level behaviours; from feeding and sexual functioning to goal-directed, flexible cognition ^4–7^. Amongst these, behavioural inhibition, memory, and aversive processing are historically considered the core, specialised functions of serotonin ^8–11^. This is underpinned by converging preclinical and human work involving *in vivo* manipulation of synaptic 5-HT, predominantly with SSRIs or depletion of its amino acid precursor tryptophan [TRP] ^7, 12^, and observing behavioural change. In humans, however, stark differences in the direction of behavioural effects are observed across similar experimental approaches ^8, 13^. For example, several studies report seemingly contradictory effects of SSRIs on tasks of aversive and reward processing (reinforcement learning); specifically, different reports show that SSRIs increase reward sensitivity ^13^, increase loss sensitivity and decreased reward sensitivity ^14^, and decrease sensitivity to both reinforcement valences ^15^. Inconsistent behavioural effects of SSRIs are also observed across other domains, including behavioural inhibition and memory processing ^12, 16–22;^ in some cases, these behavioural changes align with those seen after TRP depletion (*e.g.* reduced cognitive flexibility) despite the expectation that they would have opposing effects on net synaptic 5-HT ^16, 23^.

Determining a causal link between increased synaptic 5-HT and behaviour in humans via SSRIs is difficult due to the complex effects of SSRIs on 5-HT and co-localised neurotransmitter systems. For example, negative signalling feedback along the serotoninergic pathway following autoreceptor activation early in treatment can limit cell firing, and therefore 5-HT release, in a regionally-specific manner ^24–26^. Furthermore, deactivation of 5-HT transporters results in 5-HT uptake via dopamine transporters, leading to subsequent co-release of dopamine and 5-HT ^27^. The effect of increased dopaminergic content and signalling is seen in acute and short-term SSRI administration ^28–32^, observable in striatal, prefrontal, and hippocampal structures implicated in reward processing, behavioural inhibition and memory functioning ^27, 33–35^.

Given the complex molecular and behavioural profile of SSRIs, alternative probes which increase synaptic 5-HT may help further clarify the role of 5-HT in human behaviour and cognition. One such alternative involves the use of a selective serotonin releasing agent (SSRA) (Fig 1): unlike SSRIs which increase 5-HT levels indirectly through prolonging synaptic 5-HT, SSRAs stimulate direct exocytic release of 5-HT, without broad monoaminergic efflux (as seen in non-selective 5-HT releasers, such as MDMA) ^36, 37^. While SSRIs require ongoing neural firing for vesicular release of 5-HT into the synapse, the SSRA mechanism is not firing-dependent and thus not negated by dorsal raphe autoreceptor negative feedback which delays the therapeutic onset of action of SSRIs ^38–40^.

**Fig 1.**
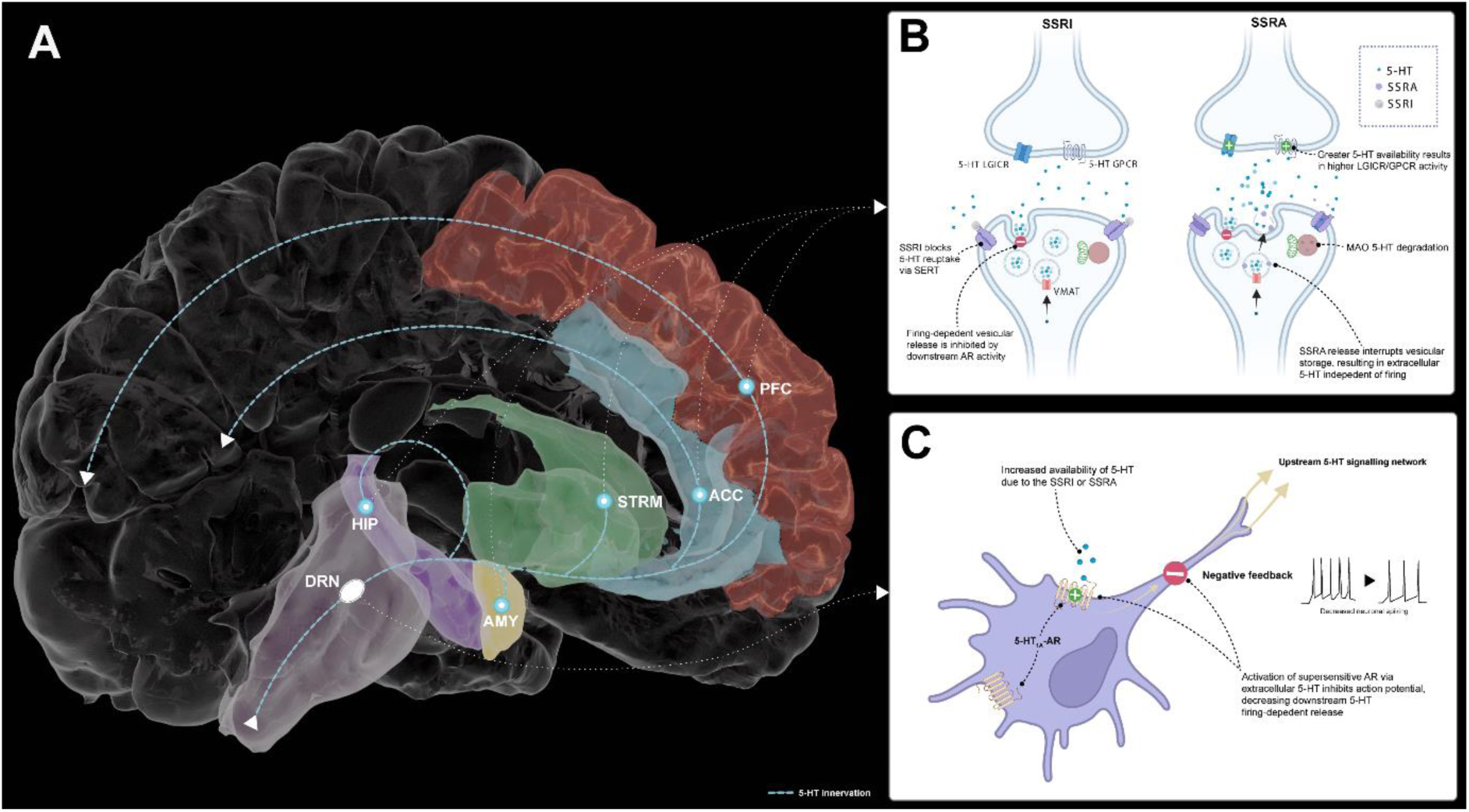
Selective serotonin releasing agent is not negated by 5-HT_1A_ supersensitivity, resulting in a rapid onset of pro-serotoninergic activity. **A.** The majority of central 5-HT innervation originates from the dorsal raphe nucleus (lilac), and is found within areas of the brain strongly implicated in mood regulation and cognitive function: amygdala (yellow), hippocampus (purple), striatal structures (green), anterior cingulate cortex (light blue) and the prefrontal cortex (red). **B.** SSRIs and SSRAs both influence extracellular presynaptic serotonin concentrations, allowing for greater serotoninergic activity, while the effects of SSRIs on synaptic 5-HT are delayed by autoreceptor hypersensitivity and may influence co-localised dopamine neurons. **C.** 5-HT_1a_ ARs are clustered in the dorsal raphe nucleus and are endogenously sensitive to extracellular serotonin, and upon activation produce a negative feedback loop which inhibits upstream firing-dependent serotonin release. **Abbreviations**: AR: autoreceptor; GPCR: G protein-coupled receptor; LGICR: Ligand-gated ion-channel receptors; MAO = Monoamine oxidase; SERT = serotonin transporter. **Note:** Original atlas meshes are credited to A. M. Winkler (Brain For Blender), which have been modified for illustrative purposes.

Until recently, it has been challenging to characterise the effects of SSRAs in humans because of the lack of available licensed pharmacological probes. However, in 2020, low dose fenfluramine (up to 26mg daily; racemic mixture) was licensed for the treatment of Dravet epilepsy ^41^. Unlike SSRIs, low dose fenfluramine directly and rapidly increases synaptic 5-HT without modifying extracellular dopamine concentration in regions involved in mood regulation such as the striatum and hippocampus ^40, 42–53^. Fenfluramine results in substantially greater extracellular 5-HT levels than the SSRI, fluoxetine, when administered at similar doses ^54^. Acute administration of fenfluramine increases synaptic 5-HT by 182-200% vs basal state ^53, 55^, while short-term administration (4-5 days) retains the increases in net 5-HT without influencing 5-HT terminal structural integrity ^53, 56^ . With its recent relicensing for epilepsy syndromes ^41^, fenfluramine provides a novel opportunity to probe the neurobehavioural effects of SSRAs in humans to answer outstanding questions about the role of synaptic 5-HT in human behaviour.

Here we use this now accessible approach to directly increase synaptic 5-HT in humans and examine its influence on domains of behaviour historically considered historically core functions of serotonin: aversive processing, behavioural inhibition, and memory. We hypothesised that the SSRA would result in a pattern of behaviour opposite to that seen with tryptophan depletion, namely reduced sensitivity to aversive outcomes, coupled with improved behavioural inhibition and memory ^16, 17, 57–62^.

## Results

A sample of 53 young, non-clinical participants were recruited (62% female; SSRA:placebo = 26:27; mean age = 20.2) and were well-matched across demographic factors (Supplementary Table 1). All participants in the final sample attended testing sessions before treatment and at follow-up (8 ± 1 day).

### Does increased synaptic serotonin change reinforcement sensitivity for reward and loss?

We investigated the effect of SSRA administration on reinforcement sensitivity for reward and loss outcomes during a probabilistic instrumental learning task described in Fig 2A ^63, 64^. During this task, participants learned the probability of outcomes associated with symbols within pairs. Each pair represented a task condition: win trials (win money or no change) and loss trials (lose money or no change). Optimal choices were made when selecting symbols which had a greater probability (70%) of leading to a favourable outcome (*i.e.,* win in win trials and no change in loss trials). Computational reinforcement learning models were fitted to participant choice during the task (see Supplementary Methods) to formalise a predicted change in optimal choice making between allocation groups. Model parameters for each trial type were derived, providing a distinct explanation of learning and decision-making behaviour throughout the task: learning rate (𝛼), explaining the rate at which outcomes modify expectations; outcome sensitivity (ρ), explaining the effective magnitude of experienced outcomes; and inverse decision temperature (𝛽), explaining the extent to which expectations inform choices (choice stochasticity). Model parameters ρ and 𝛽 were estimated across separate models. Inferential tests

**Fig 2.**
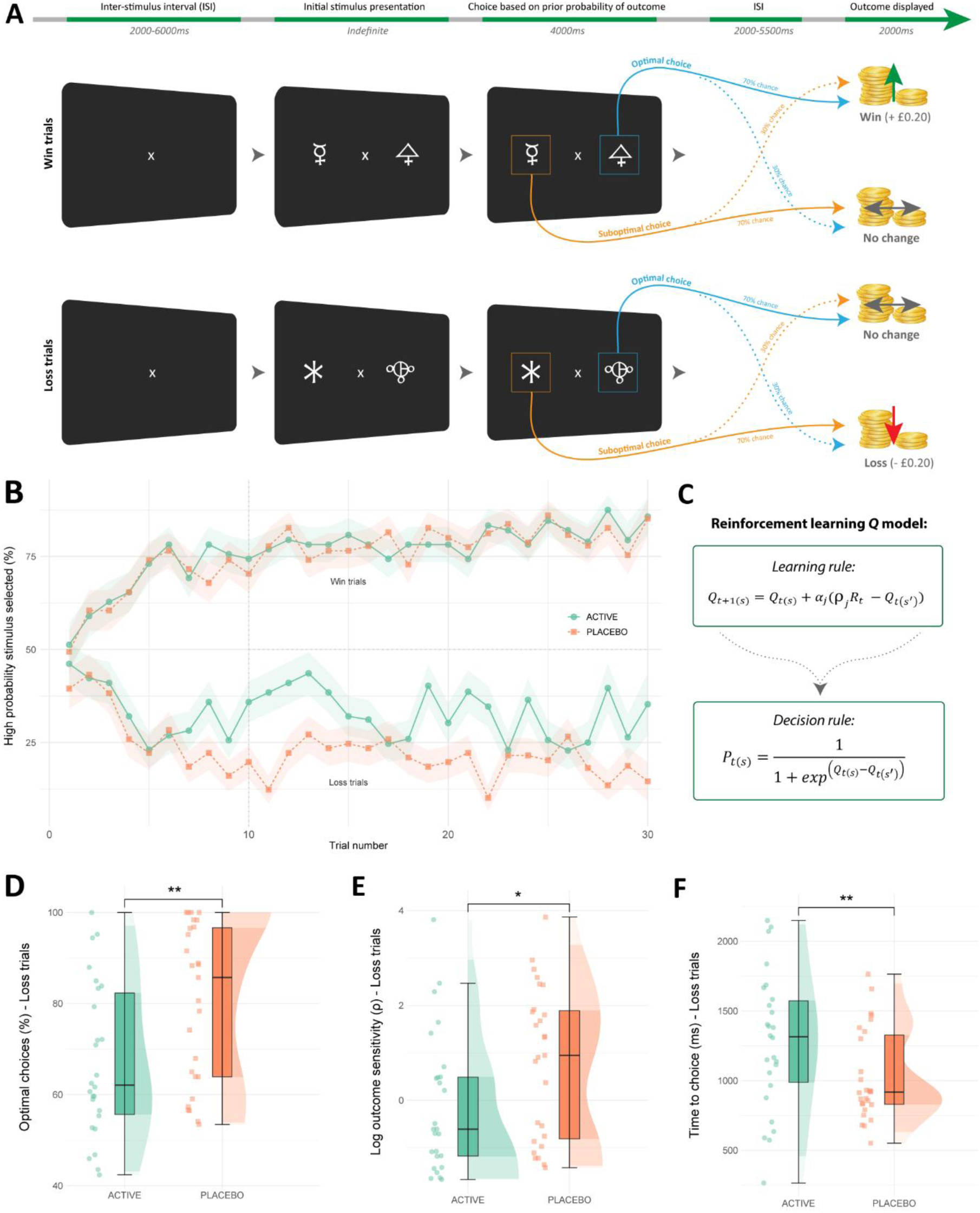
Task procedure, computational modelling, and analyses of the probabilistic instrumental learning task. **A.** Probabilistic Instrumental Learning Task flow. The task starts with a brief ISI (first screen) followed by a choice selection between one of a pair of symbols per trial (middle screen). Two novel pairs of symbols alternate throughout task blocks, with one pair representing win trials where probability of winning is higher (top row of trials) or loss trials where probability of loss is higher (bottom row of trials). Win trials result in a 20p gain or no change, while loss trials results in a 20p loss or no change. For each pair, symbols are tied to reciprocal probability values of 70% or 30%, where the outcome of a selection is displayed following each trial (final screen). Participants were instructed to select outcomes most likely to translate to maximal monetary gain which would are awarded to them at study completion. **B.** Rates of learning between allocation groups across both win and loss trials averaged across all task blocks (30 trials per trial type). ‘High probability stimulus selected’ (Y axis) is the mean percentage of choices for stimuli with a high probability of monetary win or loss. The shaded area for each line represents standard error. **C.** The *Q* computational model consists of two primary parts, a learning rule (above) which inputs to a decision rule (below). The learning rule describes how value expectation (‘𝑄_𝑡(𝑠)_’) and observed outcome (‘𝑅_𝑡_’) update on a trial-by-trial basis, where choice probability is determined via the decision rule. Model parameters alter distinct aspects of the decision-making process: outcome sensitivity (‘ρ’) and learning rate (‘𝛼′) (for further details, see Supplementary Methods). **D.** Decreased optimal choice selection in the SSRA group during loss trials. **E.** Computational modelling analysis: decreased outcome sensitivity (ρ) in the SSRA group during loss trials **F.** Increased response time in the SSRA group during loss trials only. **Note:** All panels include data for *N*=53 individuals. Error bars depict standard mean error, and half-violin plots depict the data distribution; group difference by EMM: ** *p* ≤ 0.01, * *p* ≤ 0.05.

In line with our hypothesis, SSRA allocation reduced the number of optimal choices during loss but not win trials (ANCOVA group x task condition: F[1,50] = 5.14, *p* = 0.03, η ^2^ = 0.07 [95% CI 0.00, 0.24]; loss condition EMM ± SE = -8.62 ± 3.18, *p* < 0.01, Cohen’s *d* = -0.75 [95% CI -1.30, -0.19]; reward condition EMM = 0.68 ± 3.18, *p* = 0.83) (Fig 2B-C). Consistent with this, learning models fit to the data revealed SSRA allocation reduced outcome sensitivity for loss trials only (ANCOVA group x task condition: F[1,50] = 5.73, *p* = 0.02, η ^2^ = 0.10 [0.00, 0.28]; loss condition EMM = -0.90 ± 0.43, *p* = 0.04, *d* = -0.57 [-1.11, -0.03]; reward condition EMM = 0.10 ± 0.43, *p* = 0.82) (Fig 2D). In contrast, modelled learning rate for both conditions did not vary across groups (ANCOVA group x task condition: F[1,50] = 1.22, *p* = 0.27; main effect of group: F[1,50] = 0.92, *p* = 0.34) (Supplementary Fig 2). SSRA allocation increased time to choice selection during loss conditions only (ANCOVA group x task condition: F[1,50] = 5.52, *p* = 0.02, η ^2^ = 0.11 [0.00, 0.29]; loss condition EMM = 246.0 ± 95.6, *p* = 0.01, *d* = 0.71 [0.15, 1.26]; reward condition EMM = 13.9 ± 95.6, *p* = 0.89) (Fig 2E), which would also be consistent with a relative reduction in loss sensitivity in this group.

Overall, these findings demonstrate that net increases in synaptic 5-HT (via SSRAs) decreases reinforcement sensitivity to loss outcomes while reward remains unchanged, opposite to the effect of 5-HT depletion (TRP) where loss sensitivity increases ^61, 62^. While alternative computational accounts for the observed behaviour could include increased value decay or choice stochasticity, there are no reports of 5-HT manipulation influencing these components of behaviour ^64^.

### Do SSRAs modulate behavioural inhibition, choice impulsivity, and vulnerability to aversive emotional interference?

Next, we assessed the impact of SSRA allocation on response inhibition (an index of behavioural inhibition), choice impulsivity, and interference during the Affective Interference Go/No-Go task. In this task, participants respond (‘go’) or withhold responses (‘no-go’) according to rules which change over time (*e.g.,* “do not press the button if you see a blue/yellow image”) while being exposed to emotional distractors (fearful or happy faces, or control images) (Fig 3A). SSRA allocation increased response inhibition, measured by mean percentage of accurately withheld responses to ‘no-go’ trials (ANCOVA main effect of group: F[1,47] = 11.26, *p* < 0.01, η ^2^ = 0.15 [0.00, 0.37]; all conditions EEM = 9.69 ± 2.63, *p* < 0.001, *d* = 0.60 [0.27, 0.93]) (Fig 3B). Further, groups did not differ in total accuracy for trials where a response was required (‘go’ trials) (ANCOVA main effect of group: F[1,47] = 0.83, *p* = 0.37) (Supplementary Fig 3B).

**Fig 3.**
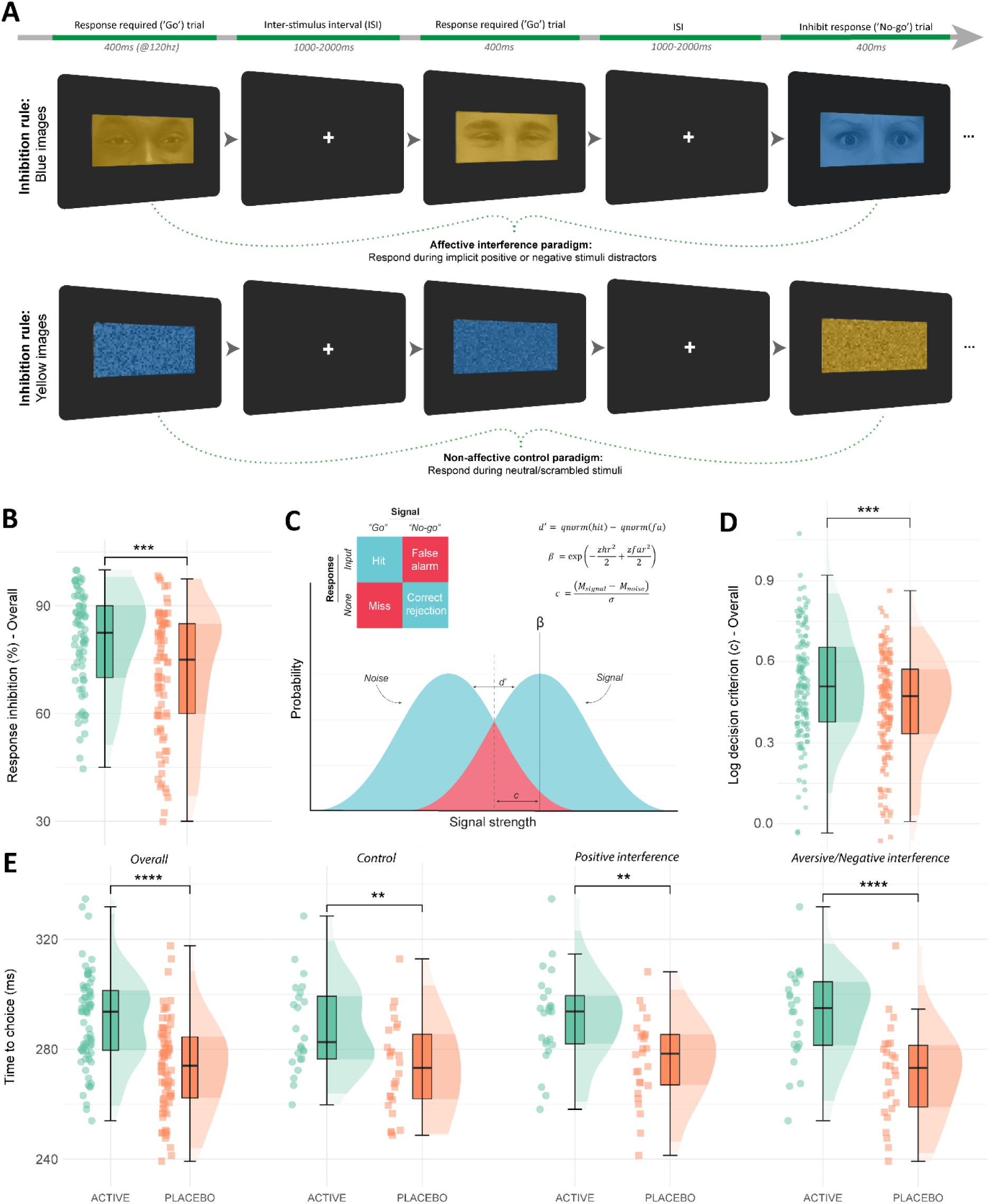
Task procedure and non-model analyses for the Affective Interference Go/No-Go task. **A.** An example of trial flow across two blocks from the affective interference go/no-go task (above), with one block during the affective interference condition and the other during the non-emotional (scrambled) control condition (below). The sequence of trials is left to right. The first two trials in each condition illustrate ‘go’ trials where participants respond with a key input (80% of trials); the third trial in the sequence illustrates a ‘no-go’ trial where participants must withhold responses (20% of trials). **B.** Higher response inhibition (mean %) performance was observed in the SSRA group compared with the placebo group across all conditions. **C.** Application of signal detection theory indices to go/no-go task, where correct and incorrect go/no-go responses are described on a sensory continuum of ‘noise’ and ‘signal’ (more details in Supplementary Materials). **D.** SSRA allocation resulted in higher values for signal detection theory criterion index ‘*c*’ (indicative of more conservative/cautious decision-making) across all task conditions. **E.** General decreases in choice impulsivity (or, choice time for correct ‘go’ trials) were observed in the SSRA group; this effect was most pronounced during aversive interference. **Note:** All panels include data for *N*=50 individuals. Error bars on each boxplot depict standard mean error, and half-violin plots depict the data distribution; group difference by EMM: **** *p* ≤ 0.0001, *** *p* ≤ 0.001, ** *p* ≤ 0.01.

Signal detection theory analyses was undertaken to determine if group differences in response inhibition were driven by perceptual decision-making (Fig 3C). SSRA allocation resulted in more cautious decision-making throughout (log criterion *c*; ANCOVA main effect of group: F[1,47] = 13.54, *p* < 0.001, η ^2^ = 0.19 [0.02, 0.39]; all conditions EMM = 0.08 ± 0.02, *p* < 0.001, *d =* 0.39 [0.16, 0.62]) (Fig 3D), but similar signal discriminability (see Supplementary Results).

SSRA allocation also resulted in reductions in choice impulsivity, indicated by increased time to choice for ‘go’ trials, across all task conditions (ANCOVA main effect of group: F[1,47] = 22.00, *p* < 0.001; η_p_^2^ = 0.27 [0.07, 0.46]) (Fig 3E). Moreover, there was an interaction between group and task interference (happy, fearful or control distractors) on choice impulsivity (ANCOVA group x task condition: F[2,95] = 3.22, *p* = 0.05, η_p_^2^ = 0.08 [0.00, 0.20]). Specifically, choice impulsivity in the SSRA group was most reduced when aversive emotional distractors were present (EMM = 21.3 ± 4.71, *p* < 0.0001, *d* = 1.28 [0.70, 1.86]) compared with both control (EMM = 14.6 ± 4.71, *p* < 0.01, *d* = 0.88 [0.31, 1.44]) and positive emotional distractors (EMM = 15.4 ± 4.71, *p* < 0.01, *d* = 0.93 [0.36, 1.50]).

Computational drift diffusion modelling (Fig 4A) was undertaken to investigate evidence accumulation patterns throughout the Affective Interference Go/No-Go task (see Supplementary Methods). SSRA allocation shifted initial choice bias (*a*z*) toward impulse control (‘no-go’, lower boundary) during aversive interference only (ANCOVA group x task condition: F[1,95] = 3.46, *p* = 0.03, η ^2^ = 0.06 [0.00, 0.17]; aversive interference EMM = -0.33 ± 0.15, *p* = 0.03, *d* = -0.60 [-1.17, - 0.04]; positive interference EMM = -0.16 ± 0.15, *p* = 0.31; control condition EMM = -0.01 ± 0.15, *p* = 0.96) (Fig 4B). Groups did not differ across other model parameters, including boundary separation (*a*) and drift rate (*v*) (see Supplementary Results). As 75% of task trials fit to the DDM were ‘go’ trials, and there was no group difference on accuracy for these trials, group differences in model parameters may not occur when accuracy is similar despite differences in choice time ^65^.

**Fig 4.**
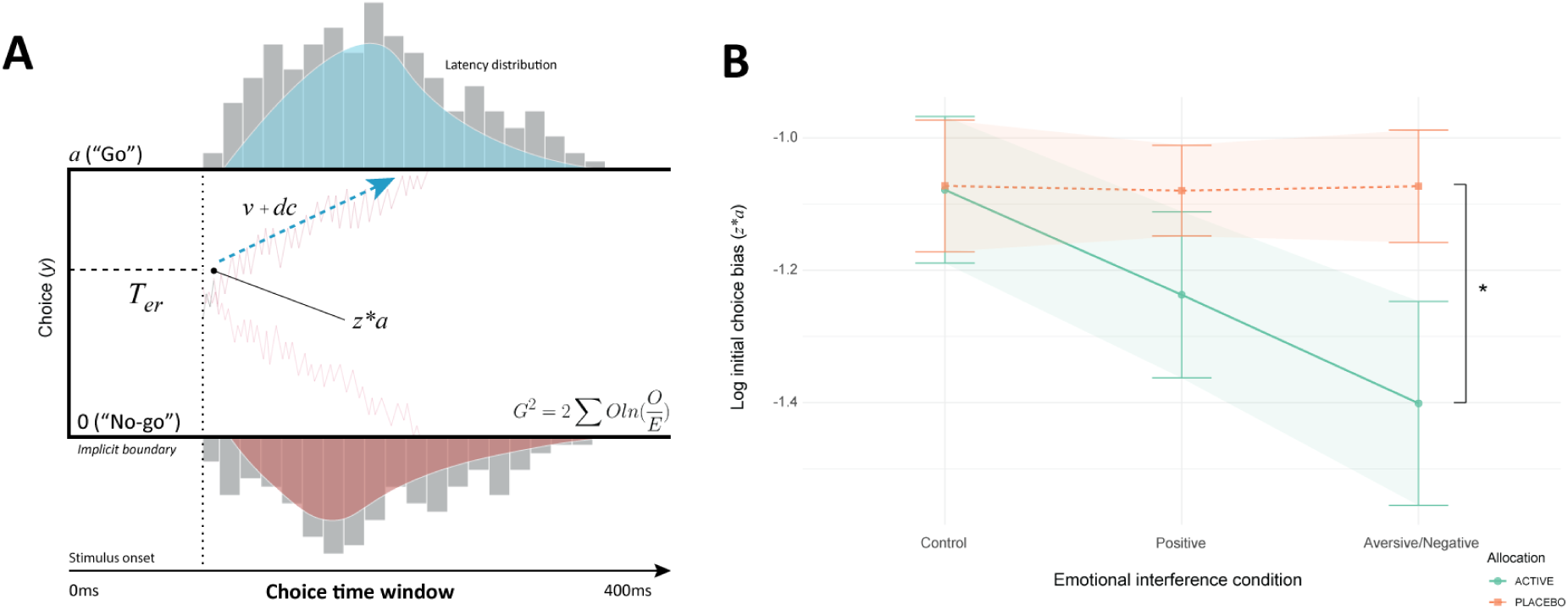
Computational drift diffusion modelling and choice bias during affective interference. **A.** The drift diffusion model describes the process of evidence accumulation and integration during the Affective Go/No-Go Task. The model was fit to observed behaviour using the Gsquare (G^2^) approach which uses maximum likelihood estimation, where choice time distributions for ‘go’ trials were divided into five quantiles: 10^th^, 30^th^, 50^th^, 70^th^ and 90^th 66, 67^. The model describes behaviour using five parameters: 1) Boundary separation (*a*), which describes the required quantity of evidence for decision-making. 2) Non-decision time (*T_er_*) is the period between stimulus onset and evidence accumulation processing where foremost sensory and perceptual processes occur, notably emotional facial expression encoding ^68^. 3) Initial choice bias (*z*a*) which represents bias toward one of the choice boundaries (*a* [‘Go’] and *0* [‘No-go’]) at the start of evidence accumulation. 4) Drift rate (*v*) describes the rate of evidence accumulation before arriving at a choice boundary. 5) Drift criterion (*dc*) is a constant applied to the mean drift rate which is evidence independent. **B.** During interference with aversive emotional information (fearful faces), SSRA allocation resulted in an initial choice bias (*z*a*) toward the impulse control (‘no-go’) choice boundary (*N* = 50). This corresponds with an increase in choice time for ‘go’ trials specifically during aversive interference in the SSRA group. **Note:** Error bars and shaded areas around each plot line depict standard mean error; group difference by EMM: * *p* ≤ 0.05.

Taken together, these findings suggest that increasing synaptic 5-HT results in a generalised enhancement of behavioural inhibition. This effect was driven specifically by more cautious decision-making, and not differences in signal discriminability or evidence accumulation rate. Moreover, increased 5-HT levels appear to shift bias towards impulse control during aversive affective interference at the start of evidence accumulation, consequentially lowering choice impulsivity.

### Assessing the influence of increased synaptic serotonin on memory processing

Finally, we assessed the influence of SSRA administration on memory function. During a task of verbal working memory processing (Verbal *n*-back; Fig 5A), participants were required to recall if a target letter occurred within a pre-specified sequential pattern (i.e., 0-, 1-, 2-, or 3-back letters ago).

**Fig 5.**
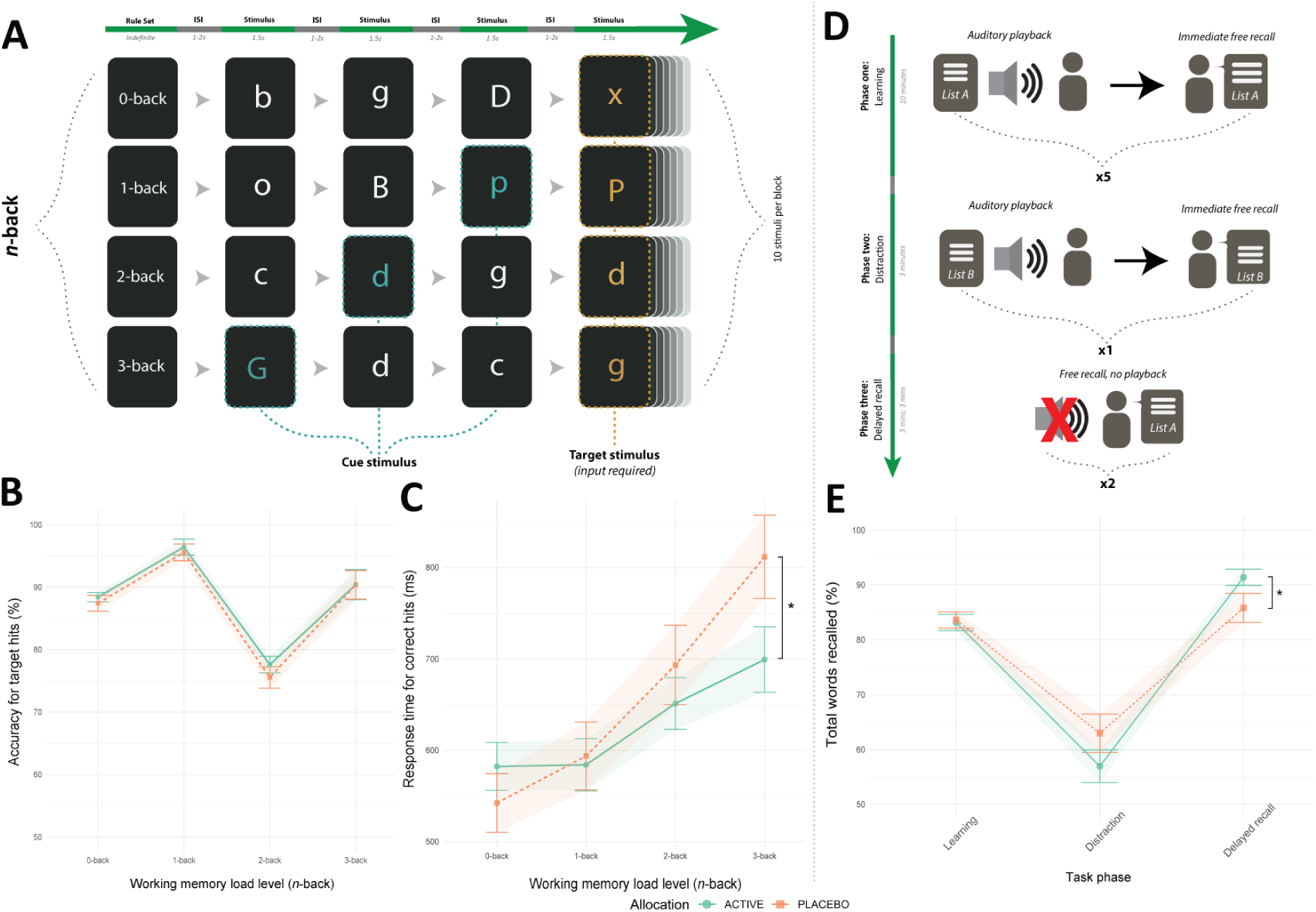
Effects of the SSRA across tasks of memory function (*n*-Back and Auditory Verbal Learning Task). **A.** Verbal *n*-back task example task flow for all four task conditions (top to bottom: 0-back, 1-back, 2-back and 3-back). The sequence of trials is left to right. Before each block of 10 stimuli, participants were given a rule for targets (*e.g.*, press spacebar if you see the same letter that appeared two letters ago [2-back]). Each condition was repeated four times (16 blocks total). **B.** No difference in target accuracy was observed across groups, while there was a significant main effect of *n*-back load on accuracy of target hits (Supplementary Results, *N* = 52)**. C.** Reduced response time for correct choices (hits) in the SSRA group at the highest load of working memory load complexity in the *n*-Back task (*N* = 52). **D.** Auditory Verbal Learning Task flow across three task phases: phase one (learning/encoding), phase two (distraction), and phase three (delayed recall). During phase one, participants listened to a recording of 15 verbal items (List A) at a slowed pace (1s gap between words), followed by an immediate free recall of list items. After this occurred five times, phase two (distraction) required learning a novel list of items (List B). Phase three (delayed recall) required free recall (without list playback) of items from List A immediately after phase two and then fifteen minutes later. **E.** The SSRA group showed increased accuracy during the delayed recall phase of the Auditory Verbal Learning Task relative to placebo (*N* = 51). **Note**: Error bars and shaded areas around each plot line depict standard mean error; group difference by EMM: * *p* ≤ 0.05.

Groups did not differ in total number of correctly recalled targets (ANCOVA group analysis: F[1,49] = 0.58, *p* = 0.45) (Fig 5B). However, during the highest task difficulty (3-back) SSRA allocation resulted in faster recall for correct targets (ANCOVA group x task condition: F[3,149] = 3.69, *p* = 0.01, η ^2^ = 0.05 [0.00, 0.13]; 3-back EMM: -112.03 ± 50.54, *p* = 0.03, *d* = -0.62 [-1.17, -0.67]) (Fig 5C).

During a task of long-term memory encoding and retrieval (Auditory Verbal Learning Task; Fig 5D), participants were required to learn a list of 15 verbal items and correctly recall these items during learning (immediate recall) and after a short period (delayed recall). SSRA allocation resulted in higher total accuracy during delayed recall but not immediate recall of learned verbal information (ANCOVA group x task condition: F[2,1474] = 6.23, *p* = 0.01, η ^2^ = 0.11 [0.08, 0.14]; delayed recall EMM = 0.84 ± 0.35, *p* = 0.02, *d* = 0.34 [0.06, 0.61]; immediate recall EMM = -0.07 ± 0.14, *p* = 0.63; distractor recall EMM = -0.90 ± 0.70, *p* = 0.20) (Fig 5E). Groups did not differ in frequency of recall repetitions or intrusions (Supplementary Figs 5–7).

Groups did not differ in terms of performance on tasks of visuo-spatial working memory (Oxford Memory Task) and implicit visual learning (Contextual Cueing Task) (see Supplementary Results, Supplementary Table 6 and Supplementary Fig 4).

Taken as a whole, these findings suggest increasing synaptic 5-HT enhances memory processing for verbal, but not visuospatial, information.

### Effects of SSRA on cortisol levels and self-report questionnaire measures

Group allocation was not related to changes in salivary cortisol concentration or self-report ratings of subjective cognition, side effects, motivation and affect (see Supplementary Results for further details). These results partly rule out the potential of motivation or affect to indirectly drive change in task behaviour ^21^.

## Discussion

The present findings demonstrate the direct effects of increased synaptic 5-HT on human behaviour, underlining its critical role in guiding decision-making across aversive and more neutral contexts. Specifically, we observed reduced sensitivity specifically for outcomes in aversive but not appetitive contexts; enhanced behavioural inhibition and increased bias favouring impulse control during aversive affective interference; and enhanced memory function for verbally-encoded information. These findings offer broad implications for longstanding theories of how central 5-HT influences human behaviour and contributes to psychiatric aetiology.

### Implications for theory of central serotonin: dichotomy of aversive and reward processing in instrumental learning

Throughout instrumental learning, increased synaptic 5-HT (via the SSRA) reduced sensitivity to aversive, but not reward-related, outcomes. This effect is opposite to that described following central depletion of serotonin with tryptophan-depletion, where enhanced negative prediction errors during probabilistic instrumental learning and bias toward aversive but not rewarding stimuli during Pavlovian conditioning have been observed ^60, 62, 69, 70^. Further, in a Pavlovian-to-instrumental transfer paradigm, independent depletion of 5-HT and dopamine respectively enhanced aversive and decreased rewarding Pavlovian-to-instrumental transfer ^71^. As SSRAs and TRP result in opposite effects on net synaptic 5-HT, the opposite behavioural pattern observed here is consistent with a key role for serotonin in modulating loss sensitivity ^38, 57^.

An absence of change in reward sensitivity from the SSRA contrasts with the effects of SSRIs in humans. Despite the shared purpose of increasing synaptic 5-HT, SSRI administration has been associated with decreased sensitivity for rewarding outcomes ^14, 15, 72^. Reduced reward sensitivity has been attributed to unwanted SSRI treatment effects, notably emotional blunting and reduced efficacy in targeting anhedonia ^73^. Importantly, SSRI administration results in indirect modulation of dopaminergic signalling pathways involved in reward processing ^28–32^. However, the SSRA used here (low dose fenfluramine, racemic mixture) retains selectivity for 5-HT ^45, 55, 74, 75^, and is inactive at dopaminergic synapses ^56, 76^, in addition to a binding affinity for 5-HT transporters which is <0.5% of that typically seen in SSRIs such as citalopram ^77^ (see the Supplementary Discussion for further details on the pharmacodynamic properties of the experimental probe and its past uses). Thus, these results highlight potentially specific effects of serotonin on loss processing, whereas contradictory effects of SSRIs previously reported may relate to effects beyond the serotonin system.

The effect of increased synaptic 5-HT on aversive but not reward processing is further supported by a body of preclinical literature. Pharmacological (fenfluramine) and optogenetic stimulation of serotonergic neurons in the dorsal raphe nucleus [DRN] results in no changes in reward processing in animal models; however, stimulation of non-serotonergic DRN neurons via amphetamine and optogenetics results in marked increases in reward processing ^78^. Moreover, increased firing of amygdala 5-HT neurons is observed during aversive but not reward prediction errors, an effect which appears to be modulated by a functionally discrete DRN to basal amygdala 5-HT pathway ^79, 80^. Accordingly, direct 5-HT depletion in amygdala and orbitofrontal cortex modulates learning about aversive but not rewarding feedback ^81^.

### A step toward uncovering the shared role of serotonin in inhibition and aversive processing

Increasing synaptic 5-HT (via the SSRA) enhanced behavioural inhibition, an effect driven by more cautious decision-making. Impairment of 5-HT function decreasing behavioural inhibition is well-observed in animals, and to lesser extent in humans ^9, 82^. However, the opposite approach of increasing synaptic 5-HT with SSRIs yields a comparably less clear picture cross-species. In humans, SSRI challenge results in improvement or no change in action cancellation ability (stop signal) ^16, 83^, while action restraint ability (go/no-go) remains unchanged or impaired ^17–19, 84^. Frontal functional activity increases during action restraint following SSRI challenge, however this is not linked to a corresponding change in ability ^18, 84^. Likewise, SSRIs yield no clear effect on behavioural inhibition in animals ^82, 85^. The seemingly irreconcilable effects of SSRIs on behavioural inhibition may be attributed to the vulnerability of the agent to experimental noise; notably, its acute-to-chronic mechanistic shift and off-target dopaminergic effects. Nevertheless, the present study is the first to demonstrate objective improvements in action restraint by increasing synaptic 5-HT. Given disorders of behavioural control and impulsivity (*e.g.,* ADHD) are associated with 5-HT dysregulation ^85^, exploring potential clinical applications of SSRAs within these populations may prove beneficial.

During behavioural inhibition, increased synaptic 5-HT resulted in a bias for impulse control during aversive interference, alongside a corresponding drop in choice impulsivity. These findings align with the longstanding conceptualisation of 5-HT as an inhibitor which becomes active in aversive contexts ^86, 87^. Indeed, in individuals with depression and tryptophan-depleted healthy adults, choice impulsivity increases for explicit negative emotional targets in a go/no-go paradigm ^88–90^. However, the effects of increased 5-HT on behavioural inhibition reported here were not experimentally confined to aversive contexts; notably, we observed a decrease in choice impulsivity during a control condition without affective interference. Potentially then, 5-HT performs an active role of limiting impulsive action more generally, but this is amplified in aversive contexts.

### Direct increases in synaptic serotonin enhance verbal memory processing

The SSRA enhanced retrieval and speed of processing during memory tasks involving verbal, but not visuospatial, information. Observable changes in memory consolidation are reliably observed following TRP depletion ^57^. SSRI challenge, however, leads to highly variable effects on long-term and episodic memory function; while improvements have been observed, typically null findings are reported ^21, 22, 91^. Unlike the SSRA, the threshold of synaptic 5-HT required for observable change may not be achieved during the brief SSRI regimen of most studies (≤ 7 days), where the problem of autoreceptor supersensitivity persists ^39, 92^. Importantly, 5-HT receptor subtypes strongly associated with memory functioning (*i.e.,* 5-HT_3,4,6_ receptors) have significantly lower binding affinities for endogenous 5-HT relative to other 5-HT receptors (*e.g*., 5-HT_1A,B,D,E,F_; 5-HT2_A-C_) ^93–96^. Thus, crossing a putative 5-HT concentration threshold may be required to observe change in memory function, potentially explaining our findings.

## Conclusion

Here we demonstrate direct effects of increased synaptic serotonin on human behaviour, underlining its critical role in guiding decision-making within aversive and more neutral contexts. In aversive contexts, increased synaptic serotonin appears to reduce sensitivity for loss outcomes, and promotes a bias toward impulse control during behavioural inhibition. In neutral contexts, increased synaptic serotonin appears to enhance behavioural inhibition by promoting cautious decisions, as well as enhancing memory recall for verbal information.

Not only do the present findings offer broad implications for longstanding theories of central serotonin, but they also demonstrate the promise of the SSRA as an experimental probe, furthering the scope of fundamental work which aims to characterise the involvement of serotonin in human behaviour, and its contribution to psychiatric aetiology in clinical samples.

Given the prominence of impaired cognition and aversive/negative emotional biases as transdiagnostic targets within psychiatry (*e.g.,* unipolar and bipolar depression; schizophrenia) ^21, 73^, investigating the therapeutic potential of the SSRA in clinical populations may be worthwhile. Such investigations may allow greater targeting of specific neurocognitive mechanisms across disorders in the absence of widespread, and often unwanted, effects including emotional blunting.

## Methods

### Participants and design

Fifty-six participants (28:28, SSRA:placebo; mean age = 20.2) were randomised to take part in the study. Recruitment occurred between June 2021 and June 2022. Potential participants were screened to exclude those who had recently used recreational drugs (3-month wash-out, except MDMA which had a wash out period of ≥ 1 year) or who were pregnant, trying to become pregnant, or who were currently breastfeeding. All participants had a BMI between 18–30 and were fluent speakers of English. For full exclusion and inclusion criteria, please see Supplementary Methods. For full details of the recruitment process, see the study CONSORT flow diagram (Fig 6).

**Fig 6.**
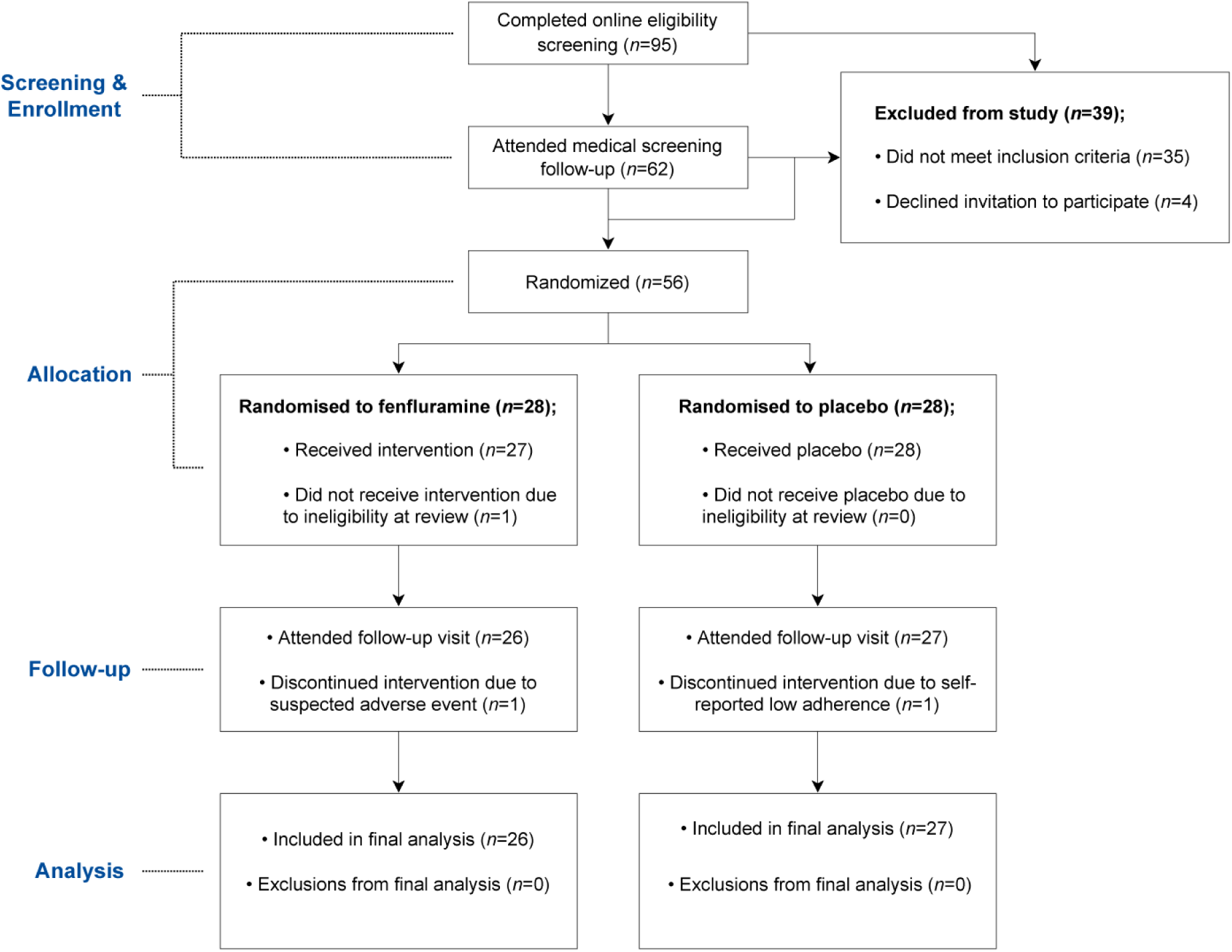
**CONSORT Diagram of participant flow throughout the study.**

Eligible participants were randomised to administration of SSRA fenfluramine hydrochloride (15mg b.i.d.; racemic mixture) or placebo for short-term administration (7-9 days), in a double-blind design. Both the SSRA and the placebo were administered orally in a flavoured aqueous solution, with the placebo lacking an active pharmaceutical ingredient. Randomisation was performed by the Clinical Pharmacy Support Unit, Oxford Health NHS Foundation Trust (Oxfordshire, United Kingdom) using a stratified block randomisation algorithm, with stratification for gender and task stimulus version (for further details on task stimulus version, see Supplementary Methods).

The study was approved by the University of Oxford Central University Research Ethics Committee (MSD-IDREC reference R69642/RE004) and pre-registered on the National Institute of Health Clinical Trials Database (NCT05026398). Prior to study participation, participants provided informed consent. All study visits were conducted at the Department of Psychiatry, University of Oxford.

### Procedure

Participants undertook two screening visits to assess study eligibility. In the first session, medical history and current medication use was assessed and the Structured Clinical Interview for DSM-V was conducted to screen for current or past psychiatric illness. In the second session, cardiovascular health (blood pressure; electrocardiography), renal and liver health (liver function, urea, and electrolyte blood tests) were assessed, and drug and pregnancy urine tests were performed. Eligible participants attended two study visits, baseline and post-intervention occurring 7, 8 or 9 days after baseline. This study period was scheduled to avoid the premenstrual week for female participants. At baseline, participants completed a battery of cognitive and emotional computer tasks and questionnaires (described in the Materials section below). Participants were then given their first dose of the SSRA or placebo and monitored for three hours during which regular blood pressure and observational checks were made. To determine cortisol levels, saliva samples were collected immediately before initial dose, one hour post-dose, and three hours post-dose. Saliva samples were immunoassayed for cortisol levels over linear calibration curves (for further details, see Supplementary Materials). After the initial dose visit, participants were asked to independently take the SSRA or placebo daily, in addition to completing daily questionnaires (see Questionnaires Measures section). At the post-intervention visit, participants completed the same task and questionnaire battery as at baseline and were then requested to estimate their allocation prior to debriefing.

### Questionnaire measures

At each study visit, participants completed self-report questionnaires measuring affect, mood, anxiety, subjective cognitive functioning, and side-effects; the Spielberger State-Trait Anxiety Inventory [STAI-T], Beck Depression Inventory II [BDI], Positive and Negative Affect Schedule [PANAS], Visual Analogue Scale [VAS], Perceived Deficit Questionnaire – Depression [PDQ-D], and side effects profile questionnaire. Participants completed the VAS and side effects questionnaires once per day between the baseline and post-intervention visits.

### Cognitive and Emotional Task Battery

Participants undertook an extensive cognitive and emotional task battery at both the initial dose visit (baseline) and follow-up visit. Participants undertook the following tasks in order: 1) Auditory Verbal Learning Task (Fig 5D) – a measure of episodic memory encoding and retrieval where accuracy of recall was the measured outcome; 2) Affective Interference Go/No-Go Task (Fig 3A) – a measure of behavioural inhibition under affective interference (positive [happy faces], aversive/negative [fearful faces], and neutral distractors) where accuracy of inhibited response to ‘no-go’ trials (response inhibition), accuracy and response time to ‘go’ trials (an index of impulsivity ^97^) were the non-model outcome measures. The block design of the task allows for analysis of set-shifting effects (executive shifting for task condition rule changes) on accuracy and response time. Participant task data was fit to a computational drift diffusion model (see Supplementary Materials for further details) which provided the following model parameters: boundary separation, initial choice bias, non-decision time, drift rate and drift criterion; 3) Verbal *N*-Back task (Fig 5A) – a measure of complex verbal working memory where accuracy and response time to ‘target’ letters (*i.e.,* matching a letter which appeared *n*-back [0, 1, 2, or 3] trials ago) were the outcome measures; 4) Probabilistic instrumental learning task ([Fig 2A] adapted from ^63^) – a measure of reward and loss sensitivity during instrumental learning, which produced non-model outcome measures which were fit to computational reinforcement learning model. Non-model outcomes were optimal choice outcome (*i.e.,* selecting the stimulus with a higher probability of a favourable outcome under each task condition: wins during win trials win or no changes during loss trials, and response time.

Computational model parameters were outcome sensitivity, learning rate and inverse decision temperature (see Supplementary Methods for further details); 5) Oxford Memory Task – a measure of visuospatial working memory which included localisation speed and stimulus selection accuracy outcomes. 6) Contextual cueing task – a measure of implicit learning and visual search ability where the outcome measure was accuracy and response times under novel/implicit cueing conditions. Full details of tasks included in this battery are included in the Supplementary Methods.

### Statistical Analysis

Data pre-processing and statistical analyses were carried out using R Software (version 4.3.1), and computational modelling was undertaken using MATLAB (R2022a) and Python (version 3.8.8). Homogeneity in demographic variables across allocation groups was assessed using chi-squared independence tests (categorical, binary variables) and two-tailed independent t-tests (continuous, discrete variables). The effect of the SSRA on outcomes across the task battery and questionnaire ratings was analysed using between-groups (SSRA vs. placebo) type III mixed model ANCOVA models on post-intervention data, with baseline performance serving as a regressor and participant as a random effect where appropriate. The approach of using baseline score as a regressor in this manner was selected as this yields greater statistical efficiency and avoids conditional bias from baseline imbalance compared with repeated-measures ANOVA ^98^ and other baseline-adjustment techniques (*e.g.,* change score between post and pre-intervention) ^99^. Post-hoc comparisons were carried out on outcome measures collected at follow-up using two-tailed estimated marginal means tests, where estimates are reported alongside standard means error; family-wise error was adjusted for via Bonferroni-Holm procedure. Effect sizes matrices are reported for the ANCOVA (partial eta squared, η ^2^) and EMM (Cohen’s *d, d*) alongside corresponding 95% confidence intervals (for effect size calculations, see Supplementary Methods). In addition to ANCOVA analysis of questionnaire data at follow-up, daily questionnaire data (VAS and side effects profile) was joined longitudinally with initial dose and follow-up visit data and analysed using linear mixed effects models with restricted maximum likelihood estimation with participant as a random effect. Salivary cortisol was analysed across three timepoints (before dose, 1- and 3-hr post-dose) using mixed linear effects modelling using time-by-allocation as an interaction term. Analyses of cortisol and self-report questionnaires are included in the Supplementary Results. All inferential analyses were carried out at the 0.05 alpha level, and significance values below 0.05 were rounded to two decimal places.

### Data availability

Source data (including raw and modelled datasets) generated for this study have been deposited on GitHub and are openly accessible here: https://github.com/mjcolwell/SSRA_human_behaviour_data_and_scripts.

### Code availability

The code used to undertake data preprocessing, modelling and inferential analyses is stored on GitHub along with the associated data: https://github.com/mjcolwell/SSRA_human_behaviour_data_and_scripts. R markdown files have been included alongside each dataset to reproduce the results reported in the present study.

## Acknowledgments

We thank Dr Sandra Tamm, Dr Angharad de Cates, and Dr Alexander Smith for their assistance in medical screening and diagnostics procedures. We thank Prof Valerie Voon for suggestions for data analysis. We thank Dr Margarita Chibalina for their assistance in biological sample handling and processing. We thank Dr Jan Willem de Gee for publishing openly available computational modelling scripts. We thank Tara Pusinelli for assistance with data collection and entry. We thank Sorcha Hamilton for help in verifying code reproducibility.

## Contributions

M.C., S.M., C.H. and P.C. designed the study. M.C., H.T., H.C. and C.W. undertook data collection. M.C., H.T., H.C. and M.B. produced preprocessing task scripts. M.C. and C.W. undertook biological specimen processing. F.S. and M.B. provided computational modelling support. M.C. undertook all computational modelling and inferential analyses. M.C., S.M., C.H., P.C., and M.B. undertook data interpretation. M.C. drafted the article and produced illustrations. All authors contributed to revisions and approval of the final draft.

## Competing Interests statement

This study was funded by a grant from Zogenix International Ltd., prior to merge with UCB Pharma, and supported by the NIHR Oxford Health Biomedical Research Centre and the NIHR Oxford Cognitive Health Clinical Research Facility. The views expressed are those of the authors and not necessarily those of the NHS, the NIHR or the Department of Health.

CH has received consultancy fees from P1vital Ltd., Janssen Pharmaceuticals, Sage Therapeutics, Pfizer, Zogenix, Compass Pathways, and Lundbeck. SM has received consultancy fees from Zogenix, Sumitomo Dainippon Pharma, P1vital Ltd., UCB Pharma and Janssen Pharmaceuticals. CH and SM hold grant income from Zogenix, UCB Pharma, Syndesi and Janssen Pharmaceuticals. CH and PC hold grant income from a collaborative research project with Pfizer. The other authors report no conflicts of interest. MB has received travel expenses from Lundbeck for attending conferences, and has acted as a consultant for J&J, Novartis, Boehringher and CHDR. He previously owned shares in P1vital Ltd.

## Notes

### Summary of Updates

-Fixed minor typo on line 282.

