## Supplementary Materials for "Direct serotonin release in humans shapes decision computations within aversive environments"

### Supplementary Methods

#### Eligibility Criteria

**Inclusion criteria:**

- Participant is willing and able to give informed consent for participation in the research
- Not currently taking any medications (except the contraceptive pill)
- Aged 18-22 years
- Male or female
- Sufficiently fluent English to understand and complete the task
- Body Mass Index above 18-30
- Weight of 40-75kg

**Exclusion criteria:**

- Current pregnancy (as determined by urine pregnancy test taken during Screening and First Dose Visit) or breast feeding
- Any past or current Axis 1 DSM-V psychiatric disorder
- Clinically significant abnormal values for liver function tests, clinical chemistry, urine drug screen, blood pressure measurement and ECG. A participant with a clinical abnormality or parameters outside the reference range for the population being studied may be included only if the Investigator considers that the finding is unlikely to introduce additional risk factors and will not interfere with the study procedures
- History of, or current medical conditions which, in the opinion of the investigator, may interfere with the safety of the participant or the scientific integrity of the study, including epilepsy/seizures, brain injury, hepatic or renal disease, severe gastro-intestinal problems, Central Nervous System (CNS) tumours, neurological conditions
- Current or past history of drug or alcohol dependency
- Use of recreational drugs (e.g. cannabis, cocaine, amphetamines) within past 3 months
- Participation in a study which uses the same computer tasks as those in the present study (determined by asking participants about previous studies participated in during screening)
- Participation in a study that involves the use of a medication within the last three months
- Smoking > 5 cigarettes per day
- Typically drinks > 6 caffeinated drinks per day (e.g. tea, coffee, coca cola, Red Bull)
- Participant is unlikely to comply with the clinical study protocol or is unsuitable for any other reason, in the opinion of the Investigator
- Current or recent use (≤1 year) of 3,4-Methylenedioxymethamphetamine (MDMA)

#### Saliva sample collection and processing

In order to determine cortisol levels, saliva samples were collected from participants immediately before taking the initial dose of the drug or placebo (baseline), one hour post-dose, and three hours post-dose. Saliva samples were collected via Salivette® Cortisol synthetic swabs. Within a 2-3 hours of collecting samples, the samples were transported to a laboratory where they were rendered acellular by centrifugation upon receipt. Prior to immunoassay, all samples were stored in the Neurosciences Building (Department of Psychiatry) in a de-identified form at -30°C. When the randomisation code was broken, all samples were immunoassayed over linear calibration curves via a Salimetrics salivary cortisol ELISA kit (#1-3002) according to the manufacturer protocol [1]. Cortisol concentration is expressed as a measure of µg/dL. Salivary cortisol was analysed across three timepoints (before dose, 1-hr and 3-hr post-dose) using mixed linear effects modelling using time-by-allocation as an interaction term.

#### Cognitive and Emotional Task Battery

The Probabilistic Instrumental Learning task (Version 220916, adapted from [2]) measures reward and loss sensitivity following instrumental learning (Fig 2A). This task requires participants to select between one of a pair of symbols per trial; two novel pairs of symbols alternate throughout task blocks, with one pair representing a high-probability-win condition (outcomes: 20p gain or no change) or a high-probability-loss condition (outcomes: 20p loss or no change). For each pair, symbols are tied to reciprocal probability values of 70% (gain/loss) or 30% (no change), where the outcome of selection is displayed following each trial. Participants were instructed to select outcomes most likely to translate to maximal monetary gain which would are awarded to them at study completion. Variables which task records includes response time, optimal selection per trial paradigm, and total gained. Per previous implementations of the task, only the last 40 trials per block (20 high-probability loss, 20 high-probability win) were analysed given that this is where the learning plateau typically occurs [2,3] Using non-model task data, learning curves were generated as a temporal-learning representation of optimal selection across each trial paradigm per block (Fig 1B).

The Affective Go/No-Go Task [AGNG] (Version 1.2) [4] measures executive control and impulsivity in under conditions of affective interference (Fig 3A). During the affective interference condition, blue and yellow rectangles are superimposed on emotional distractor images which were pairs of eyes expressing 'fear' or 'happy' emotions. The non-affective control condition displays blue and yellow rectangles superimposed on scrambled versions of the emotional distractor images with matched cropping and luminance. The task is balanced across three block types depending on the condition during the inhibitory stimuli (*i.e.,* 'fearful distractor', 'happy distractor' and 'no distractor/scramble'). Before each block new rules are given for participants (*e.g.,* "If you see a blue image, do not press spacebar"). Inhibitory ('no-go') and input required ('go') stimuli are displayed at a 1:3 ratio, respectively. Stimuli are displayed for 400ms with interstimulus intervals displayed at a pseudorandom jitter between 1000-2000ms. The task was designed for and displayed on a high refresh rate monitor (1080p, 120hz) for greater frame time accuracy/synchronisation.

The *N*-back task is a task of verbal working memory (Version 1.2) [5], wherein a sequence of letters is displayed centre-screen for an interval of 1000ms per letter. Participants are prompted before each sequence to indicate if a letter corresponded to a previously presented letter that occurred n-trials ago (one trial ago [1-back], two trials ago [2-back], three trials ago [3-back], or if the trial contained the letter 'x' [0-back]). Errors of commission and omission are recorded alongside response time.

The Contextual Cueing Task (CCT; adapted from [6]) measures implicit learning and visual search ability. During the task participants are to identify the orientation of a target stimulus which was onscreen among flanker stimuli. Half of trials include a contextual cueing element where the stimulus array was identical as the subsequent trial, while the other half contains novel arrays. In line with previous analyses of the task [6], accuracy difference scores (cued – novel trials) were analysed, in addition to response time differences (cued – novel trials). These variables were analysed via mixed ANCOVA modelling with the task stage (first half [blocks 1 & 2] or second half [blocks 3 & 4]) serving as a within-subjects factor.

The Rey Auditory Verbal Learning Task (RAVLT) [7] measures episodic memory encoding, recall and retrieval. In this implementation of the task, participants are played an audio recording of 15 commonplace nouns (List A) and prompted to recall them once the recording ended. This process is repeated five times before a novel list of commonplace nouns (List B) was played and immediate recall of the novel list was prompted. Participants are then asked to free recall List A items (short delay), and then repeat this process after a twenty minute period (long delay). Throughout the task, the number of items correctly recalled, the number of repeated recalls (repetitions) and the number of incorrect items (intrusions) are measured.

The short-form Oxford Memory Test (OMT; adapted from [8]) measures visuospatial complex working memory. During the task, 1 or 3 fractals are displayed on the screen for a brief period of 1 or 3 seconds, followed by a 1 or 4 second interstimulus period. Following this, participants are presented with two stimuli (one presented in the subsequent array and one flanker), where participants are to recall the previously presented stimuli and then drag it to its location where it was onscreen prior to the interstimulus period. Accuracy and response time for identification and localisation are collected for this task.

Participants completed all tasks in the task battery at the initial dose and follow-up visits; participants tasks in the same order each visit: 1) AVLT, 2) AGNG, 3) N-Back, 4) PILT, 5) OMT, and 6) OMT. In the randomisation algorithm, one of the two strata (alongside gender) was task stimulus version. Task stimulus version refers to versions of tasks across the cognitive and emotional task battery in which the stimuli presentation was varied to deter practice effects. There were two task stimulus versions (1 and 2) which, depending on randomisation, would be administered at the initial dose visit or follow-up visit, accordingly. All tasks within the battery were designed to have two stimulus versions except the Oxford Memory Task and Contextual Cueing Task.

#### Effects Size Calculations

Effect sizes were calculated for both mixed-effects ANCOVA models and EMM models. For the ANCOVA models, partial eta squared ($\eta_p^2$) was calculated by applying *eta_squared* function (from the 'effectsize' R package) to the mixed effect model. For EMM comparisons, cohen's *d* was calculated by applying the *eff_size* function (from the 'emmeans' R package) to an EMM derived from the outcome variable at follow-up.

#### Computational Modelling – Probabilistic Instrumental Learning Task

Data from the probabilistic instrumental learning task were fit to computational reinforcement learning models used previously [9,10], based on the $Q$ model described by Pessiglione *et al.* (2006). For the outcome sensitivity model, the value of selecting a stimulus was updated on a trial-by-trial basis in the model using the equation below, where the value of the non-selected stimulus was reciprocally updated:

$$Q_{t+1(s)} = Q_{t(s)} + \alpha_j \left( \rho_j R_t - Q_{t(s')} \right)$$

The learning expectation $Q_{t(s)}$ refers to the value attached to a stimulus/symbol ($s$) at a given time/trial ($t$), while $R_t$ refers to the positive (coded as 1 for win in win trials or no change in loss trials) or negative (coded as 0 for no change in win trials or loss in loss trials) outcome observed. Learning rate $\alpha_j$ and outcome sensitivity $\rho_j$ parameters are set for each trial type $j$ (win or loss trial). As per previous work [10], the learning expectation at initial state $Q_{0(s)}$ was set to 0.5. The unchosen option $Q_{t(s')}$ was not kept constant but rather updated based on the reciprocal outcome. $Q$ values of the first two paired stimuli were aggregated to provide a value of choice probability via the following softmax function:

$$P_{t(s)} = \frac{1}{1 + exp^{\left(Q_{t(s)} - Q_{t(s')}\right)}}$$

As with outcome sensitivity ρ, separate values for inverse decision temperature $\beta$ were derived for both trial paradigm types using a similar $Q$ model approach. Model parameters were estimated by calculating posterior probability of the two trial paradigms using a 110 by 100 grid, followed by obtaining the expected value of each marginal likelihood parameter. Values for each model parameter were log transformed prior to inferential analyses. It is worth noting that parameters $\beta$ and ρ result in similar effects on choice, despite acting on separate aspects of decision and learning throughout the task. Indeed, when fitting the model to the study data, the values for $\beta$ and ρ resulted in a perfect correlation (Supplementary Fig 1). All model parameters were log transformed and then inferentially analysed using ANCOVA and EMM approaches.

#### Signal Detection Theory Indices and Computational Drift Diffusion Modelling – Affective Interference Go/No-Go Task

Signal detection theory indices were derived using behavioural data from the Affective Interference Go/No-Go task. Required variables were calculated from task data: "Hit" (Correct 'Go' response), "Miss" (Missed 'Go' response), "False alarm" (Incorrect response to 'No-go' trial), and "Correct Rejection" (Correctly missed response to 'no-go' trial). These four variables were processed through the R package '*Psycho*' [11], and produced three indices (decision criterion [*β*], sensitivity index [*d'*] and alternate decision criterion [*c*]) which were log transformed. Each index was calculated using the following algorithms [12] :

$$d' = qnorm(hit) - qnorm(fa)$$

$$\beta = \exp\left(-\frac{zhr^2}{2} + \frac{zfar^2}{2}\right)$$

$$c = \frac{\left(M_{signal} - M_{noise}\right)}{\sigma}$$

While signal detection theory provides an estimate of signal discriminability between the groups, we decided to also fit observed behaviour to drift diffusion models (DDMs) to further understand behavioural patterns observed in the non-model data (i.e., differences in choice impulsivity across task conditions). The DDM provides a computational, mechanistic account of the evidence accumulation process during the task. We used a previously published, publicly available DDM approach for Go/No-Go data that relies on '*PyMC*' and '*HDDM*' Python packages [13–16], which resulted in high parameter recovery. Three models were fit to three task conditions: control (trials where no emotional distractors were present), positive interference (trials where happy distractor stimuli were present), and negative interference (trials where negative distractor stimuli were present). The model approach used assumed trials were independent of each other [14,17], and therefore the sequence of trials was assumed to be irrelevant to the evidence accumulation process. Participant data was fit using the gsquare approach which relies on maximum likelihood estimation, and is described as follows [14]:

$$G^2 = 2\,\Sigma\left(O \ln\left(\frac{O}{E}\right)\right)$$

For this approach, response time distributions for 'go' trials were divided into five quantiles: 10th, 30th, 50th, 70th and 90th. As the gsquare approach does not use hierarchical Bayesian fitting, the conditional independence assumption does not apply. The following model parameters were fit within the DDM: boundary separation (*a*), initial choice bias (y(0) = *z*a*), non-decision time ($T_{er}$), drift rate (*v*) and drift criterion constant (*dc*). Initial choice bias and drift criterion were fixed per the previous approach [13]. The model is described using the following stochastic differential equation:

$$\Delta y = s \cdot v \cdot \Delta T_{er} + dc \cdot \Delta T_{er} + N(0, c^2 \cdot \Delta T_{er})$$

All model parameters were log transformed and then inferentially analysed using ANCOVA and EMM approaches.

### Supplementary Results

#### Placebo/Drug Allocation Guess

At study completion, the placebo group (70%) were better than chance at correctly guessing their allocation compared with the SSRA (50%), however this difference was not statistically significant ($\chi^2$ = 3.92, *p* = 0.27).

#### Salivary Cortisol Analysis

Mean concentration of salivary cortisol (µg/dL) was analysed throughout the initial dose period (Supplementary Table 2). Before initial dose, salivary cortisol was at its lowest across the fenfluramine (mean = 0.12 ± 0.09) and placebo (mean = 0.13 ± 0.06) groups. One hour post-dose, salivary cortisol peaked in the fenfluramine (mean = 0.15 ± 0.10) and placebo (mean = 0.16 ± 0.12) groups. Three hours post-dose, salivary cortisol was sustained in the fenfluramine group (mean = 0.15 ± 0.09) but reduced in the placebo group (mean = 0.14 ± 0.05). In a linear mixed effects model, neither treatment allocation (β = 0.01, 95% CI [-0.03, 0.06], t(112) = 0.54, p = 0.59) nor the interaction term time-by-allocation (β = -0.01, 95% CI [-0.038, 0.018] t(104)= -0.71, *p* = 0.48) significantly explained variance in a model of change in salivary cortisol before and after initial dose.

#### Self-report measures

There was no group difference across most self-report ratings of cognition, affect and mood (Supplementary Table 3). There was a group effect on negative PANAS items (ANCOVA main effect: F[1,38] = 5.00, p = 0.03, $\eta_p^2$ = 0.12 [0.00, 0.32]), however the follow-up EMM analysis did not indicate a between-groups difference (EEM = -0.75 ± 0.48, *p* = 0.12). There was no significant time-by-group interaction in the longitudinal analysis of daily ratings of negative and positive VAS items (see Supplementary Table 5).

For the side effects profile, there was group difference for the 'loss of appetite' item which bordered significance (ANCOVA main effect: F[1,47] = 3.96, *p* = 0.052). No further group effects were seen on the other side effect items (see Supplementary Table 4), and there were no significant time-by-group interactions on daily ratings of all side effect items (see Supplementary Table 5).

#### Probabilistic Instrumental Learning Task – Inverse decision temperature analyses

As expected, there was an equivalent effect observed for the inverse temperature as with the outcome sensitivity parameter. Specifically, SSRA allocation reduced inverse decision temperature for loss trials only (ANCOVA group x task condition: F[1,50] = 5.73, $p = 0.02$, $\eta_p^2 = 0.10$ [0.00, 0.28]; loss condition EMM = -0.90 ± 0.43, $p = 0.04$, $d = -0.57$ [-1.11, -0.03]; reward condition EMM = 0.10 ± 0.43, $p = 0.82$).

#### Contextual cueing task

Baseline-adjusted ANCOVA found no significant main effect of group allocation on accuracy difference (cued – novel trials) on the contextual cueing task at the follow-up (ANCOVA: F[1,50] = 0.96, $p = 0.332$) (Supplementary Fig. 2A). Similarly, no significant main effect of group allocation was observed on response time difference (cued – novel trials) during the task (ANCOVA: F[1,50] = 0.04, $p = 0.84$), while a simple main effect of task stage (first half/second half) was observed (ANCOVA: F[1,50] = 17.10, $p < 0.001$; $\eta_p^2 = 0.26$ [0.08, 0.44]). Mean response time differences for each allocation group across novel or cued stimuli is detailed in Supplementary Fig 2B.

#### Oxford Memory Test

Baseline-adjusted ANCOVA found no significant main effect of allocation on 7 of 8 metrics of visuospatial working memory performance on the OMT at follow-up, as summarised in Supplementary Table 6. A significant main effect of group allocation on identification time at follow-up was observed (ANCOVA: F[1,47] = 5.56, $p = 0.02$; $\eta_p^2 = 0.01$ [0.00, 0.06]), however a post-hoc EMM analysis found no significant difference in identification time between allocation groups (EMM = 0.16 ± 0.09, $p = 0.09$).

#### *N*-Back Task

There was a main effect of *n*-back level (0-, 1-, 2-, or 3-back) on accuracy for targets (ANCOVA F[3,149] = 67.58, $p < 0.001$, $\eta_p^2 = 0.39$ [0.27, 0.49]) and response time (ANCOVA F[3,149] = 39.91, $p < 0.001$, $\eta_p^2 = 0.16$ [0.06, 0.26]). There was no effect of allocation on accuracy for identifying non-target distractors (F[1,49] = 2.76, $p = 0.10$).

#### Rey Auditory Verbal Learning Task – intrusions and repetitions

Baseline-adjusted ANCOVA found no significant main effect of group allocation on number of intrusions (F[1,48] = 0.26, $p = 0.61$) and repetitions (F[1,48] = 0.02, $p = 0.89$), while a significant main effect of trial type was observed on intrusions (F[2,48] = 4.38, $p = 0.01$, $\eta_p^2 = 0.01$ [0.00, 0.01]). and repetitions (F[2,48] = 4.53, $p = 0.01$, $\eta_p^2 = 0.01$ [0.00, 0.01]). Mean repetitions and intrusions across each trial type is detailed in Supplementary Figs. 4-5, respectively.

#### Affective Interference Go/No-Go Task

There was no significant interaction of group and set-shifting (rules changing or remaining the same across blocks) on response time via ANCOVA (F[1,47] = 0.03, $p = 0.86$) or response inhibition (F[1,47] = 0.06, $p = 0.81$), while there was a simple effect of set-shifting on response time only (F[1,347] = 5.18, $p = 0.02$, $\eta_p^2 = 0.02$ [0.00, 0.05]).

In the analysis of signal detection theory indices, SSRA allocation resulted in more conservative responses (bias index log β) across all task conditions (ANCOVA main effect of group: F[1,47] = 12.67, $p < 0.001$, $\eta_p^2 = 0.14$ [0.00, 0.36]; all conditions EMM = 0.24 ± 0.08, $p < 0.01$, $d = 0.35$ [0.12, 0.58])). Further, while there was a main effect of group on sensitivity index *d'* (ANCOVA main effect of group: F[1,47] = 7.01, $p = 0.01$, $\eta_p^2 = 0.08$ [0.00, 0.28]), the follow-up EMM analysis did not indicate a between-groups difference (EEM = 0.15 ± 0.08, $p = 0.06$).

From the computational drift diffusion analysis, ANCOVA modelling revealed no significant effect of group on further model parameters: drift rate ($F[1,47] = 1.01$, $p = 0.31$), boundary separation ($F[1,47] = 1.15$, $p = 0.29$), non-decision time ($F[1,47] = 0.01$, $p = 0.93$), and drift bias ($F[1,47] = 0.09$, $p = 0.76$).

#### Supplementary tables

**Supplementary Table 1. Demographic characteristics across allocation groups**

| | Fenfluramine (*n*=26) | Placebo (*n*=27) | Inferential analysis [a] |
|---|---|---|---|
| Age (Years), M (S.D.) | 20.19 (1.36) | 20.15 (1.29) | 0.82 |
| Gender, *N* male:female | 10:16 | 10:16 | 1.00 |
| Body mass index, M (S.D.) | 22.36 (3.48) | 22.75 (2.57) | 0.77 |
| Contraceptive use, yes:no | 6:8 | 9:7 | 0.65 |
| Native language, *N* | | | 0.67 |
| English | 19 | 20 | |
| Chinese | 3 | 3 | |
| Other | 4 | 3 | |
| Time in education (Years), M (S.D.) | 15.17 (1.22) | 15.27 (1.73) | 0.83 |
| Highest Educational Attainment, *N* | | | 0.79 |
| High School / Sixth form | 18 | 20 | |
| Undergraduate degree | 7 | 6 | |
| Postgraduate degree | 1 | 0 | |
| Not applicable | 0 | 1 | |
| Family History – Mental Health, yes:no | 5:21 | 8:18 | 0.49 |

[a] Values represent significance values inferential analyses. These values pertain to Welch's two Sample t-test where differences between group means were analysed, and Pearson's Chi-squared where differences between frequency/ratio distributions were analysed.

**Supplementary Table 2. Mean concentration (µg/dL) salivary cortisol throughout the initial dose period**

| | Active *(n=26)* | Placebo *(n=27)* |
|---|---|---|
| **OMT Domain** | **Mean ± SD** | **Mean ± SD** |
| *$T_0$: Before Dose* | 0.12 ± 0.06 | 0.13 ± 0.06 |
| *$T_1$: 1 hour post-dose* | 0.15 ± 0.10 | 0.16 ± 0.12 |
| *$T_2$: 3 hours post-dose* | 0.15 ± 0.09 | 0.14 ± 0.05 |

**Supplementary Table 3. Subjective outcome measures of cognition, affect, and mood across allocation groups – post-intervention descriptive statistics and inferential analysis**

| | Fenfluramine (*n*=26) M (S.D.) | Placebo (*n*=27) M (S.D.) | Inferential analysis [a] | |
|---|---|---|---|---|
| | | | *F*-statistic [df] | *p* |
| **Cognition** | | | | |
| *Perceived Deficits Questionnaire* | | | | |
| Baseline, M (S.D.) | 10.27 (11.80) | 15.12 (13.60) | -- | -- |
| Follow-up, M (S.D.) | 11.39 (13.32) | 11.37 (12.89) | 0.00 [1,50] | 0.996 |
| **Affect** | | | | |
| *Positive and Negative Affect Schedule* | | | | |
| Negative items, baseline, M (S.D.) | 10.86 (1.28) | 12.00 (1.81) | -- | **--** |
| Negative items, follow-up, M (S.D.) | 10.86 (1.24) | 11.61 (1.97) | 5.00 [1,38] | **0.0313** [b] |
| Positive items, baseline, M (S.D.) | 30.09 (6.37) | 28.17 (7.76) | -- | -- |
| Positive items, follow-up, M (S.D.) | 27.62 (6.74) | 26.22 (7.90) | 0.277 [1,38] | 0.602 |
| *Visual Analogue Scale* | | | | |
| Negative items, baseline, M (S.D.) | 148.92 (87.76) | 171.55 (68.73) | -- | -- |
| Negative items, follow-up, M (S.D.) | 152.12 (86.90) | 155.62 (79.94) | 0.29 [1,46] | 0.593 |
| Positive items, baseline, M (S.D.) | 736.13 (110.33) | 664.38 (83.23) | -- | -- |
| Positive items, follow-up, M (S.D.) | 699.62 (101.57) | 710.15 (99.07) | 0.00 [1,44] | 0.984 |
| **Mood** | | | | |
| *Beck Depression Inventory* | | | | |
| Baseline, M (S.D.) | 2.92 (3.14) | 5.00 (4.44) | -- | -- |
| Follow-up, M (S.D.) | 3.08 (2.99) | 2.60 (3.24) | 0.05 [1,43] | 0.826 |
| *Spielberger Trait Anxiety Subscale* | | | | |
| Baseline, M (S.D.) | 23.32 (5.51) | 23.09 (3.99) | -- | -- |
| Follow-up, M (S.D.) | 23.00 (5.64) | 22.38 (5.03) | 0.07 [1,36] | 0.790 |
| *Spielberger State Anxiety Subscale* | | | | |
| Baseline, M (S.D.) | 1.52 (2.00) | 2.26 (2.47) | -- | -- |
| Follow-up, M (S.D.) | 1.48 (1.83) | 2.33 (3.00) | 3.03 [1,36] | 0.090 |

[a] Inferential analysis via baseline-adjusted ANCOVA model across allocation groups (active vs placebo).

[b] Post-hoc EMM analysis revealed no score difference on negative PANAS items across allocation groups (EMM = -0.75 ± 0.50, 95% CI [-1.76, 0.26], *p* = 0.141), while at baseline placebo scored significantly higher on negative PANAS items than fenfluramine (EMM = -1.14 ± 0.46, 95% CI [-2.06, -0.21], p = 0.02, Hedges' *g* = -0.70). Between visits, the mean score for this item did not change in fenfluramine group while the placebo group reduced by a mean difference of 0.39.

**Supplementary Table 4. Side effects profile for allocation groups – post-intervention descriptive statistics and inferential analysis**

| *Side effect domain* | Fenfluramine (*n*=26) M (S.D.) | Placebo (*n*=27) M (S.D.) | Inferential analysis [a] | |
|---|---|---|---|---|
| | | | *F*-statistic [df] | *p* |
| **Appetite, decreased** | | | | |
| Baseline, M (S.D.) | 0.19 (0.40) | 0.35 (0.71) | -- | -- |
| Follow-up, M (S.D.) | 0.52 (0.82) | 0.16 (0.37) | 3.96 [1,47] | 0.052 |
| **Appetite, increased** | | | | |
| Baseline, M (S.D.) | 0.15 (0.37) | 0.17 (0.39) | -- | -- |
| Follow-up, M (S.D.) | 0.04 (0.20) | 0.12 (0.33) | 2.16 [1,47] | 0.148 |
| **Drowsiness/Fatigue** | | | | |
| Baseline, M (S.D.) | 0.42 (0.70) | 1.13 (0.82) | -- | -- |
| Follow-up, M (S.D.) | 0.68 (0.69) | 0.28 (0.54) | 3.28 [1,47] | 0.076 |
| **Insomnia** | | | | |
| Baseline, M (S.D.) | 0.00 (0.00) | 0.09 (0.29) | -- | -- |
| Follow-up, M (S.D.) | 0.12 (0.44) | 0.16 (0.37) | 0.49 [1.47] | 0.489 |
| **Sexual side effects** | | | | |
| Baseline, M (S.D.) | 0.00 (0.00) | 0.09 (0.42) | -- | -- |
| Follow-up, M (S.D.) | 0.12 (0.33) | 0.04 (0.20) | 0.21 [1,47] | 0.649 |
| **Sweating** | | | | |
| Baseline, M (S.D.) | 0.04 (0.20) | 0.09 (0.29) | -- | -- |
| Follow-up, M (S.D.) | 0.04 (0.20) | 0.00 (0.00) | 0.98 [1,47] | 0.327 |
| **Tremors** | | | | |
| Baseline, M (S.D.) | 0.00 (0.00) | 0.00 (0.00) | -- | -- |
| Follow-up, M (S.D.) | 0.00 (0.00) | 0.08 (0.28) | 2.09 [1,47] | 0.155 |
| **Agitation** | | | | |
| Baseline, M (S.D.) | 0.04 (0.20) | 0.13 (0.34) | -- | -- |
| Follow-up, M (S.D.) | 0.16 (0.47) | 0.04 (0.20) | 1.54 [1,47] | 0.221 |
| **Anxiety** | | | | |
| Baseline, M (S.D.) | 0.15 (0.37) | 0.22 (0.42) | -- | -- |
| Follow-up, M (S.D.) | 0.00 (0.00) | 0.08 (0.40) | 1.85 [1,47] | 0.180 |
| **Diarrhoea** | | | | |
| Baseline, M (S.D.) | 0.00 (0.00) | 0.00 (0.00) | -- | -- |
| Follow-up, M (S.D.) | 0.28 (0.84) | 0.04 (0.20) | 1.92 [1,47] | 0.172 |
| **Dry Mouth** | | | | |
| Baseline, M (S.D.) | 0.08 (0.27) | 0.04 (0.21) | -- | -- |
| Follow-up, M (S.D.) | 0.24 (0.44) | 0.12 (0.33) | 0.00 [1,47] | 1.000 |
| **Indigestion** | | | | |
| Baseline, M (S.D.) | 0.00 (0.00) | 0.04 (0.21) | -- | -- |
| Follow-up, M (S.D.) | 0.00 (0.00) | 0.08 (0.28) | 2.06 [1,47] | 0.158 |
| **Nausea** | | | | |
| Baseline, M (S.D.) | 0.04 (0.17) | 0.09 (0.29) | -- | -- |
| Follow-up, M (S.D.) | 0.16 (0.55) | 0.04 (0.20) | 1.02 [1,47] | 0.318 |
| **Upset stomach** | | | | |
| Baseline, M (S.D.) | 0.08 (0.27) | 0.09 (0.29) | -- | -- |
| Follow-up, M (S.D.) | 0.16 (0.55) | 0.12 (0.33) | 0.10 [1,47] | 0.760 |

[a] Inferential analysis via baseline-adjusted ANCOVA model across allocation groups (active vs placebo)

**Supplementary Table 5. Mixed-effects linear modelling of longitudinal (daily) subjective ratings data**

| | Model estimate [a] | 95% CI | *t* value | *p* |
|---|---|---|---|---|
| *Visual Analogue Scale* | | | | |
| *Positive items* | -17.63 | -67.80, 32.55 | -0.69 | 0.494 |
| *Negative items* | 28.20 | -8.32, 64.32 | 1.35 | 0.178 |
| *Side Effects Profile* | | | | |
| Appetite, decreased | -0.07 | -0.35, 0.21 | -0.50 | 0.622 |
| Appetite, increased | 0.11 | -0.02, 0.24 | 1.74 | 0.089 |
| Drowsiness/Fatigue | -0.02 | -0.23, 0.26 | -0.13 | 0.901 |
| Insomnia | 0.01 | -0.06, 0.23 | 1.18 | 0.242 |
| Sexual side effects | 0.01 | -0.12, 0.14 | 0.14 | 0.888 |
| Sweating | 0.04 | -0.01, 0.09 | 1.48 | 0.147 |
| Tremors | 0.01 | -0.03, 0.06 | 0.57 | 0.571 |
| Agitation | 0.01 | -0.11, 0.11 | 0.05 | 0.959 |
| Anxiety | 0.07 | -0.04, 0.18 | 1.27 | 0.211 |
| Diarrhoea | -0.11 | -0.26, 0.04 | -1.41 | 0.166 |
| Dry Mouth | -0.02 | -0.24, 0.21 | -0.15 | 0.879 |
| Indigestion | 0.05 | -0.01, 0.12 | 1.63 | 0.110 |
| Nausea | -0.05 | -0.20, 0.10 | -0.64 | 0.528 |
| Upset stomach | -0.05 | -0.21, 0.12 | -0.54 | 0.595 |

[a] Inferential analysis via baseline-adjusted mixed-effects linear model across allocation groups (active vs placebo).

**Supplementary Table 6. Oxford Memory Test (visual working memory) ANCOVA summary with main effect of treatment allocation**

| | Active *(n=24)* | Placebo *(n=24)* | | | |
|---|---|---|---|---|---|
| **OMT Domain** | **Mean ± SD** | **Mean ± SD** | **F-Statistic** [a] | **Degrees of freedom** | ***P*-Value** |
| *Absolute Error* | 74.33 ± 44.94 | 79.06 ± 55.36 | 0.40 | 1,47 | 0.53 |
| *Misbinding value* | 0.07 ± 0.09 | 0.09 ± 0.10 | 0.08 | 1,47 | 0.79 |
| *Guessing value* | 0.09 ± 0.08 | 0.08 ± 0.07 | 0.19 | 1,47 | 0.67 |
| *Targeting value* | 0.85 ± 0.15 | 0.85 ± 0.15 | 0.03 | 1,47 | 0.86 |
| *Identification time* | 1.64 ± 0.70 | 1.48 ± 0.61 | 5.56 | 1,47 | 0.02 |
| *Localisation time* | 3.30 ± 1.09 | 3.20 ± 1.00 | 0.43 | 1,47 | 0.52 |
| *Proportion correct* | 0.96 ± 0.05 | 0.96 ± 0.05 | 0.00 | 1,47 | 1.00 |
| *Imprecision value* | 50.67 ± 17.49 | 51.49 ± 19.44 | 0.08 | 1,47 | 0.78 |

[a] via baseline-adjusted ANCOVA type II

#### Supplementary Figures

*(One figure per page, centred.)*

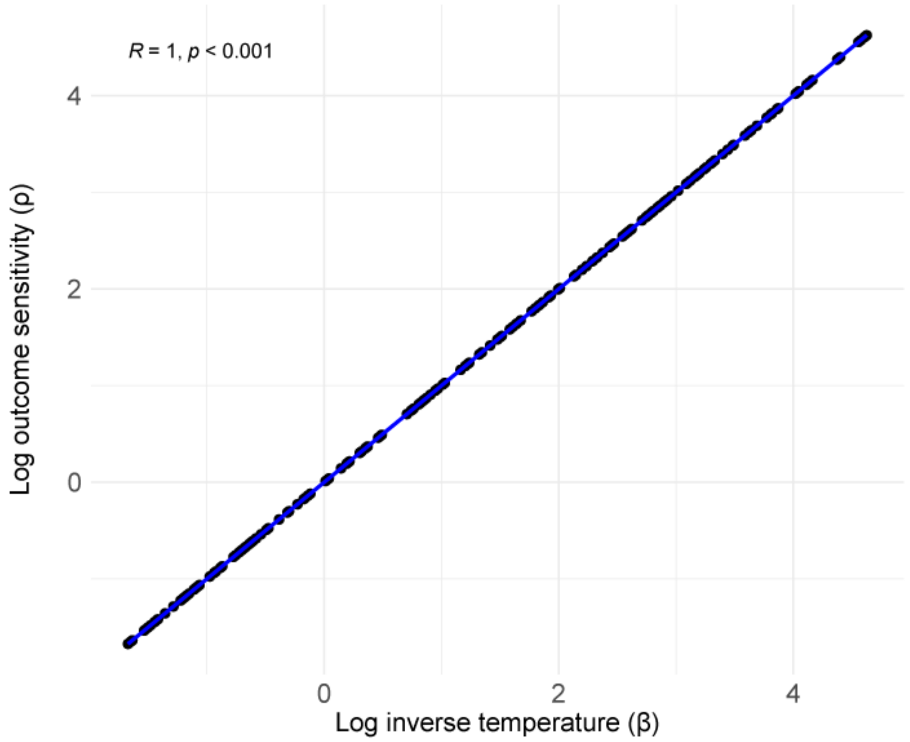

**Supplementary Fig 1**. Relationship between each reinforcement learning model parameter, outcome sensitivity ρ and inverse temperature $\beta$.
**Note:** R refers to Pearson correlation coefficient.

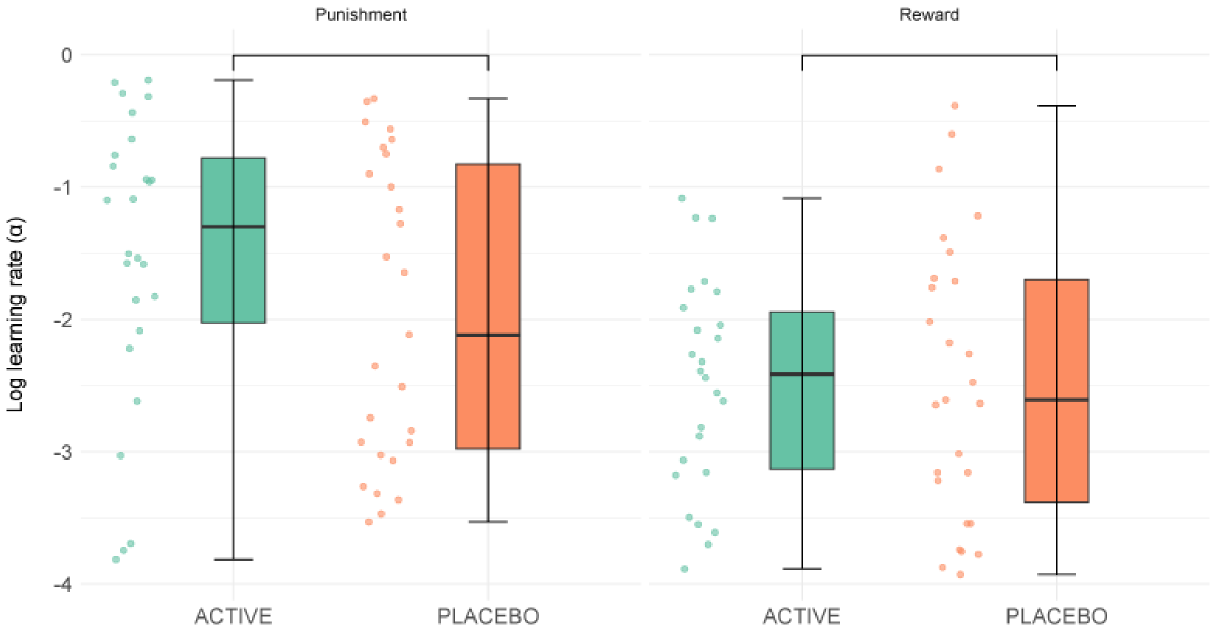

**Supplementary Fig 2.** No effect of allocation of allocation on learning rates for loss/punishment and reward trials on the Probabilistic Instrumental Learning Task

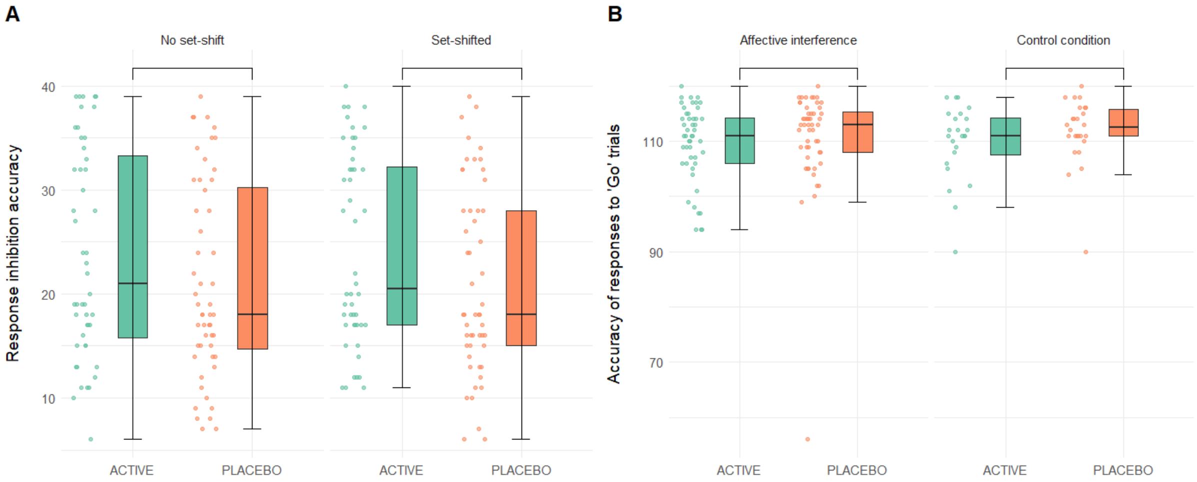

**Supplementary Fig 3. No effect of allocation (A) on response inhibition across set-shifted/non-set-shifted blocks and (B) on accuracy for 'go' trials at follow-up (Affective Go/No-go task).**

**Note:** No significant interaction between group allocation and set-shifting (rules changing or remaining the same across blocks) on response inhibition performance was observed ($F[1,48] = 0.04$, $p = 0.84$). Baseline-adjusted ANCOVA revealed no significant effects of group allocation on commission (go trial) performance ($F[1,48] = 0.01$, $p = 0.98$).

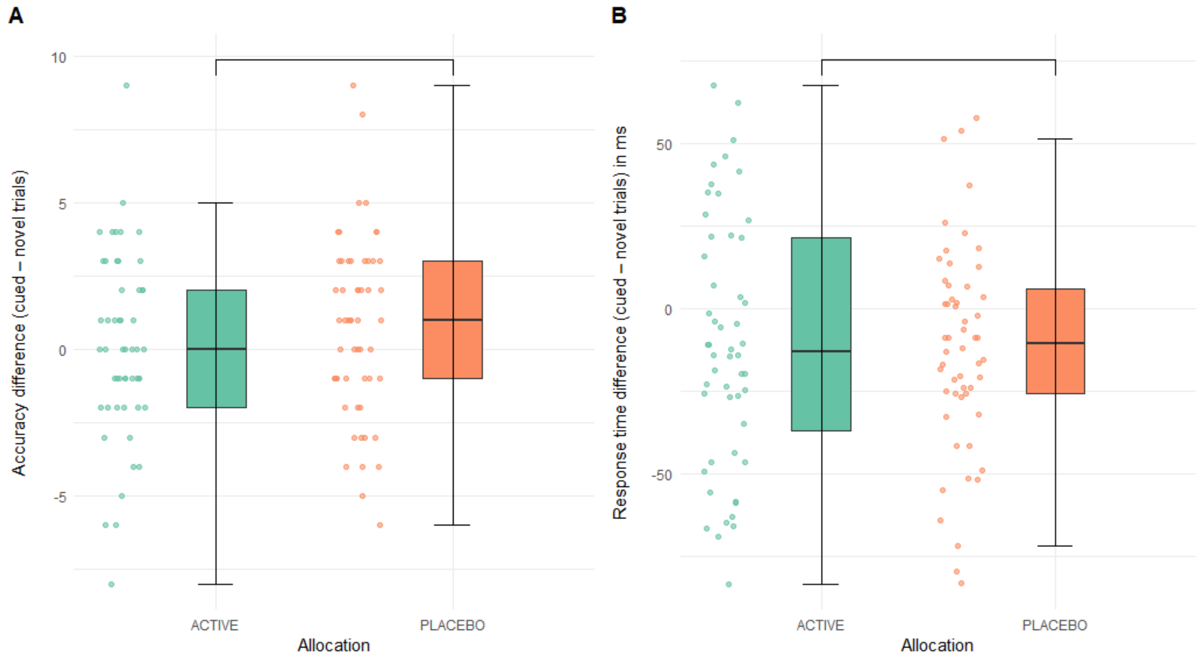

**Supplementary Fig 4. Contextual cueing task performance: no effect of allocation on (A) accuracy difference (A) (cued – novel trials correct choices) and (B) response time difference (cued – novel trials) in milliseconds (ms).**

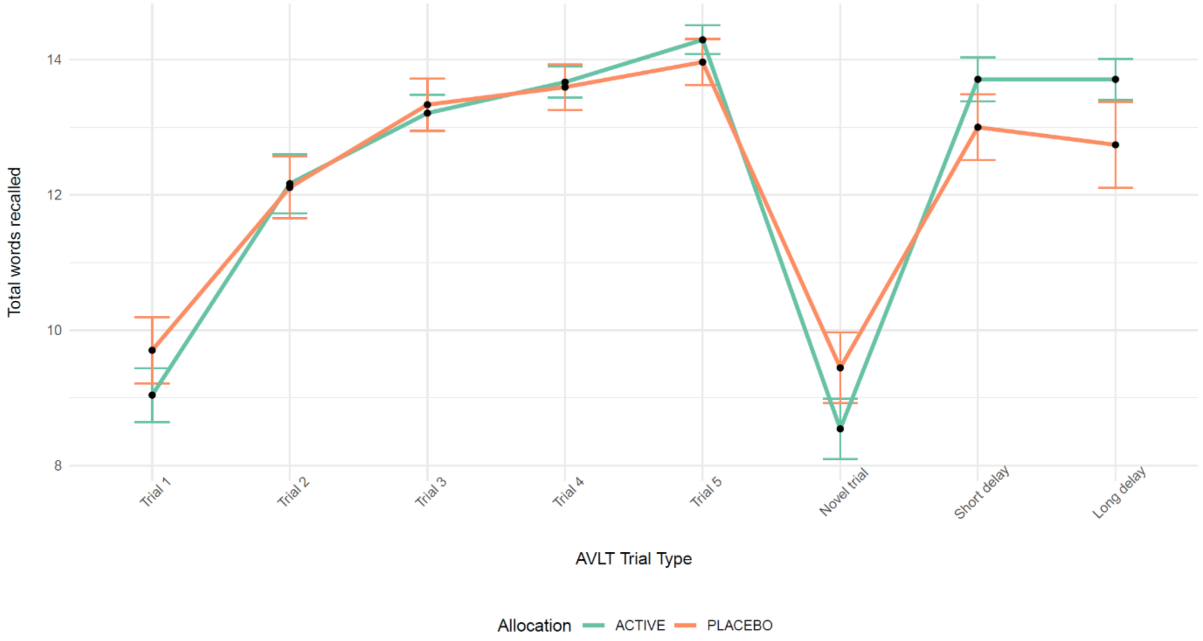

**Supplementary Fig 5. Split by allocation group, total words recalled on the AVLT at follow-up across each trial type (Learning [Trials 1 – 5], Novel trial, and Free recall (Short delay and Long delay)**

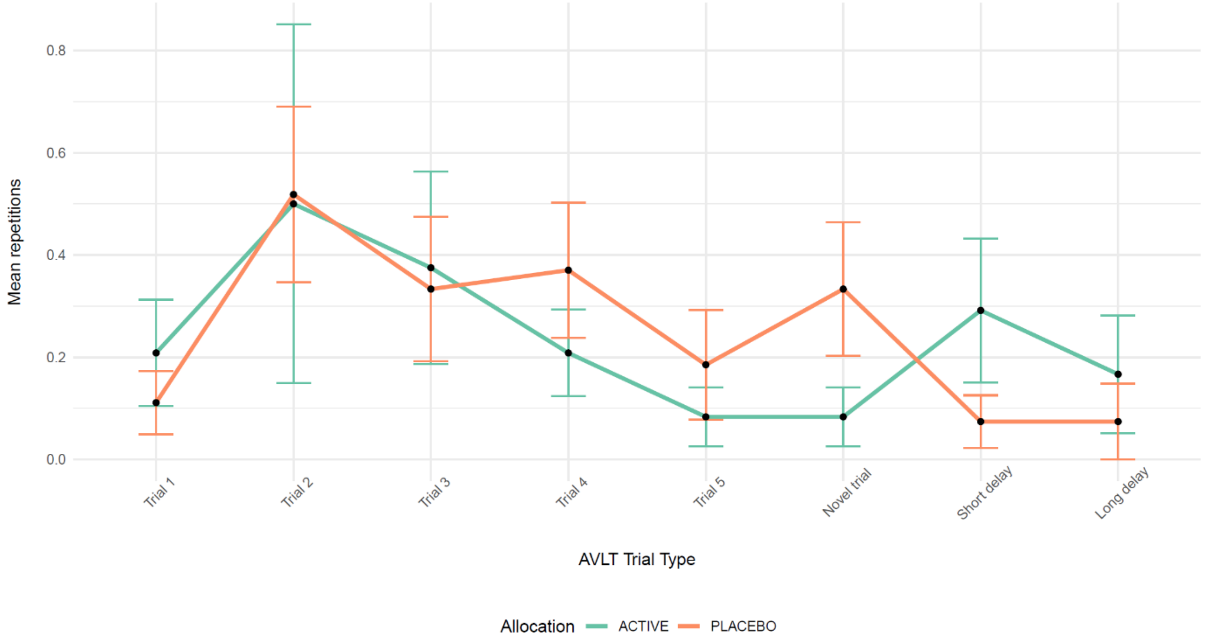

**Supplementary Fig 6. Split by allocation group, mean repetitions on the AVLT at follow-up across each trial type (Learning [Trials 1 – 5], Novel trial, and Free recall (Short delay and Long delay)**

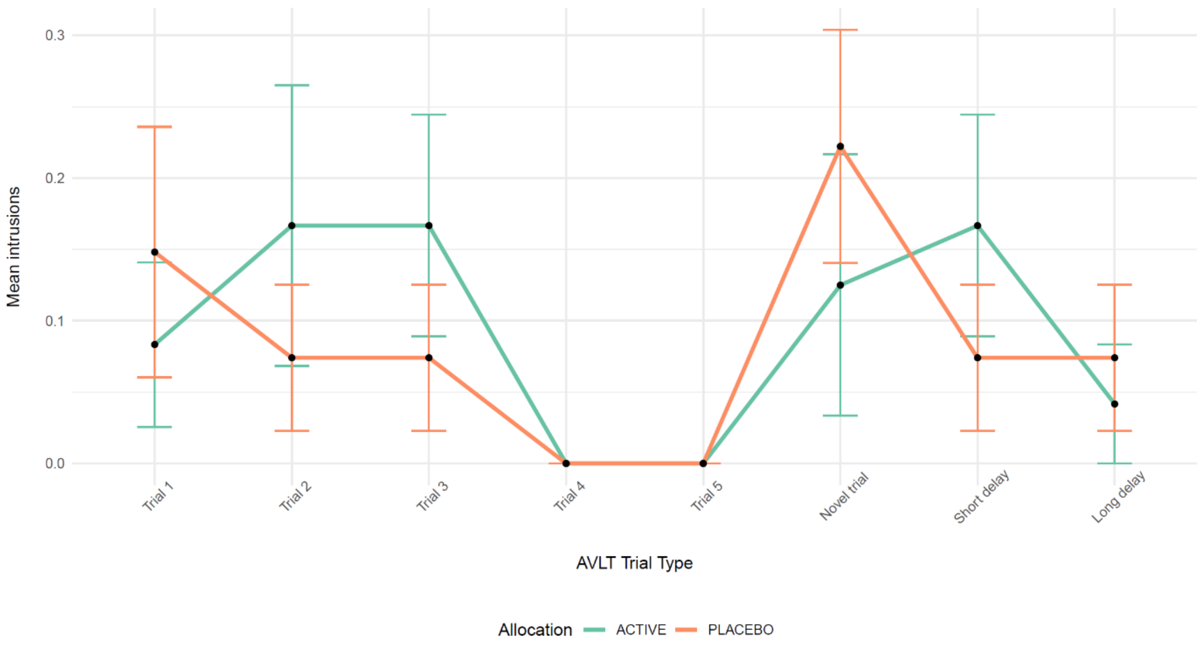

**Supplementary Fig 7. Split by allocation group, mean intrusions on the AVLT at follow-up across each trial type (Learning [Trials 1 – 5], Novel trial, and Free recall (Short delay and Long delay)**

### Supplementary Discussion

#### Auxiliary pharmacodynamic effects of SSRA *dl*-fenfluramine and its neuroactive metabolite

As with most psychoactive substances, including SSRIs [18–22], the SSRA fenfluramine has some potential 'off-target' pharmacological effects; these include activity at sigma-1 and 5-$HT_2$ receptors.

While retaining neurotransmitter selectivity for the serotoninergic system at low doses, fenfluramine has modest affinity for the 5-$HT_2$ receptor subtype and sigma-1 ($\sigma_1$) receptors. In the case of 5-HTR, these effects appear to be specific to 5-$HT_{2A/B/C}$R [23], while there is indirect/inconclusive evidence of the involvement of other receptors such as 5-$HT_4$ [24]; however, the binding affinity of fenfluramine for 5-$HT_{2A/B/C}$R is at most < 1% of that of competitive endogenous 5-HT [23,25,26]. Certainly, given high concentration of 5-HT following SSRA fenfluramine and the finite availability of 5-HT receptors [27–29], the resulting auxiliary effects of 5-HTR agonism/antagonism from fenfluramine on its broader neurobehavioral profile are likely negligible. Moreover, fenfluramine appears to be both a positive allocentric modulator and antagonist of $\sigma_1$R [30,31]; while endogenous neurosteroid sigma-1 agonists may be potentiated through this mechanism, leading to improvements in cognitive ability [31–33], it is unclear to what extent this effect may be offset by the sigma-1 antagonist properties of fenfluramine in the healthy brain [30,34–36].

Pharmacodynamic data on norfenfluramine, the neuroactive metabolite of fenfluramine, at similar doses is relatively limited; in one study, *d*-norfenfluramine administration in mice resulted in small increases (relative to 5-HT) of synaptic noradrenaline but not dopamine [37]. However, in two further studies noradrenaline and dopamine levels were unaltered by *d*-norfenfluramine and *dl*-norfenfluramine administration [38,39]. Moreover, *dl*-fenfluramine administration in humans produces plasma concentrations of *dl*-fenfluramine and *dl*-norfenfluramine at a 1:3 ratio, respectively, with only a fraction of that being *d*-norfenfluramine [37,40]. Nevertheless, contrasting the neurobehavioural profile of SSRA fenfluramine with *S*-enantiomers (selective to 5-HT) of SRAs such as 4-methyl-N-methylcathinone [41], once clinically available, would help elucidate the potential neurobehavioural contribution of the other pharmacological effects of SSRA fenfluramine.

#### Past investigations of the influence of *d-/dl*-fenfluramine on behaviour

Interpretation of our findings in the context of past work on fenfluramine is challenging. The few available neurobehavioral studies of fenfluramine in humans are limited by small and heterogeneous samples, with the most recent published almost two decades ago [42–44]. In this early work, higher doses of *dl*-fenfluramine or d-enantiomer dexfenfluramine were administered which may diminish selectivity for 5-HT [45–48] and have greater potential for neurotoxicity [49,50]. It is important, therefore, that independent attempts to replicate the present findings are undertaken to determine the reliability of low dose fenfluramine as a pro-serotonergic probe.

Our findings align with preliminary work in patients with Dravet Syndrome showed improvements in caregiver ratings of executive functioning following low dose fenfluramine administration [51] .

#### Absence of pseudo-specific effects

Research aiming to examine the effect of pharmacological drugs on non-affective cognitive processing must consider the role of pseudo-specificity (*i.e.,* indirect effects of affective processing on cognition [52]). Here we observed no group differences in self-reports of general affective functioning (*e.g.,* motivation), allowing us to rule out the potential pseudo-specific effects of the SSRA on enhancements to behavioural inhibition and memory.
